# Pervasive lesion segregation shapes cancer genome evolution

**DOI:** 10.1101/868679

**Authors:** Sarah J. Aitken, Craig J. Anderson, Frances Connor, Oriol Pich, Vasavi Sundaram, Christine Feig, Tim F. Rayner, Margus Lukk, Stuart Aitken, Juliet Luft, Elissavet Kentepozidou, Claudia Arnedo-Pac, Sjoerd Beentjes, Susan E. Davies, Ruben M. Drews, Ailith Ewing, Vera B. Kaiser, Ava Khamseh, Erika López-Arribillaga, Aisling M. Redmond, Javier Santoyo-Lopez, Inés Sentís, Lana Talmane, Andrew D. Yates, Colin A. Semple, Núria López-Bigas, Paul Flicek, Duncan T. Odom, Martin S. Taylor

## Abstract

Cancers arise through the acquisition of oncogenic mutations and grow through clonal expansion^1, 2^. Here we reveal that most mutagenic DNA lesions are not resolved as mutations within a single cell-cycle. Instead, DNA lesions segregate unrepaired into daughter cells for multiple cell generations, resulting in the chromosome-scale phasing of subsequent mutations. We characterise this process in mutagen-induced mouse liver tumours and show that DNA replication across persisting lesions can generate multiple alternative alleles in successive cell divisions, thereby increasing both multi-allelic and combinatorial genetic diversity. The phasing of lesions enables the accurate measurement of strand biased repair processes, the quantification of oncogenic selection, and the fine mapping of sister chromatid exchange events. Finally, we demonstrate that lesion segregation is a unifying property of exogenous mutagens, including UV light and chemotherapy agents in human cells and tumours, which has profound implications for the evolution and adaptation of cancer genomes.

## Main text

Sequencing and analysis of cancer genomes has identified a wealth of driver mutations and mutation signatures^1, 3^. These mutation signatures have revealed how environmental mutagens cause genetic damage and elevate cancer risk^3–5^. Analyses of the genes driving carcinogenesis has identified underlying cellular deficiencies that can help direct cancer therapies^6–8^. The diversity of mutation patterns identified from cancer genome sequencing is testament to the temporal and spatial heterogeneity of exogenous and endogenous exposures, mutational processes, and germline variation amongst patients. A recent study of diverse human cancers identified 49 distinct single base substitution signatures, with almost all tumours demonstrating evidence of at least three signatures^3^.

Such intrinsic heterogeneity leads to overlapping mutation signatures that confound our ability to accurately disentangle the biases of DNA damage and repair, or to interpret the dynamics of clonal expansion. Transcription coupled repair (TCR) partially reduces the mutational burden on the actively transcribed genome, but its analysis has been inherently limited by many factors. For example, the requirement to compile tumours of diverse aetiologies to achieve sufficient mutations to analyse, the absence of accurate measurements of transcription in the cell of origin, and an inability to evaluate on which strand mutagenic lesions occur^9^.

We reasoned that a more controlled and genetically uniform cancer model system would overcome some of these limitations and complement human cancer studies. By effectively re-running cancer evolution hundreds of times, we aimed to explore the processes of mutagenesis, DNA repair, and clonal expansion in cancer development at high resolution, and with good statistical power.

We chemically induced liver tumours in fifteen-day-old (P15) male C3H/HeOuJ inbred mice (subsequently C3H, n=104) using a single dose of diethylnitrosamine (DEN), thus greatly extending our previous study^10^. To provide a genetic comparison and a validation dataset in a divergent mouse strain^11^, we treated a cohort of CAST/EiJ mice with DEN (subsequently CAST, n=84). Tumours were isolated 25 or 38 weeks post-DEN treatment for C3H and CAST, respectively (**Fig. 1a**, **Supplementary Table 1**); each tumour was processed in parallel for histopathological and DNA analysis. Tumours fulfilling the histological inclusion criteria (**Methods**) proceeded to sequencing.

**Fig. 1.**
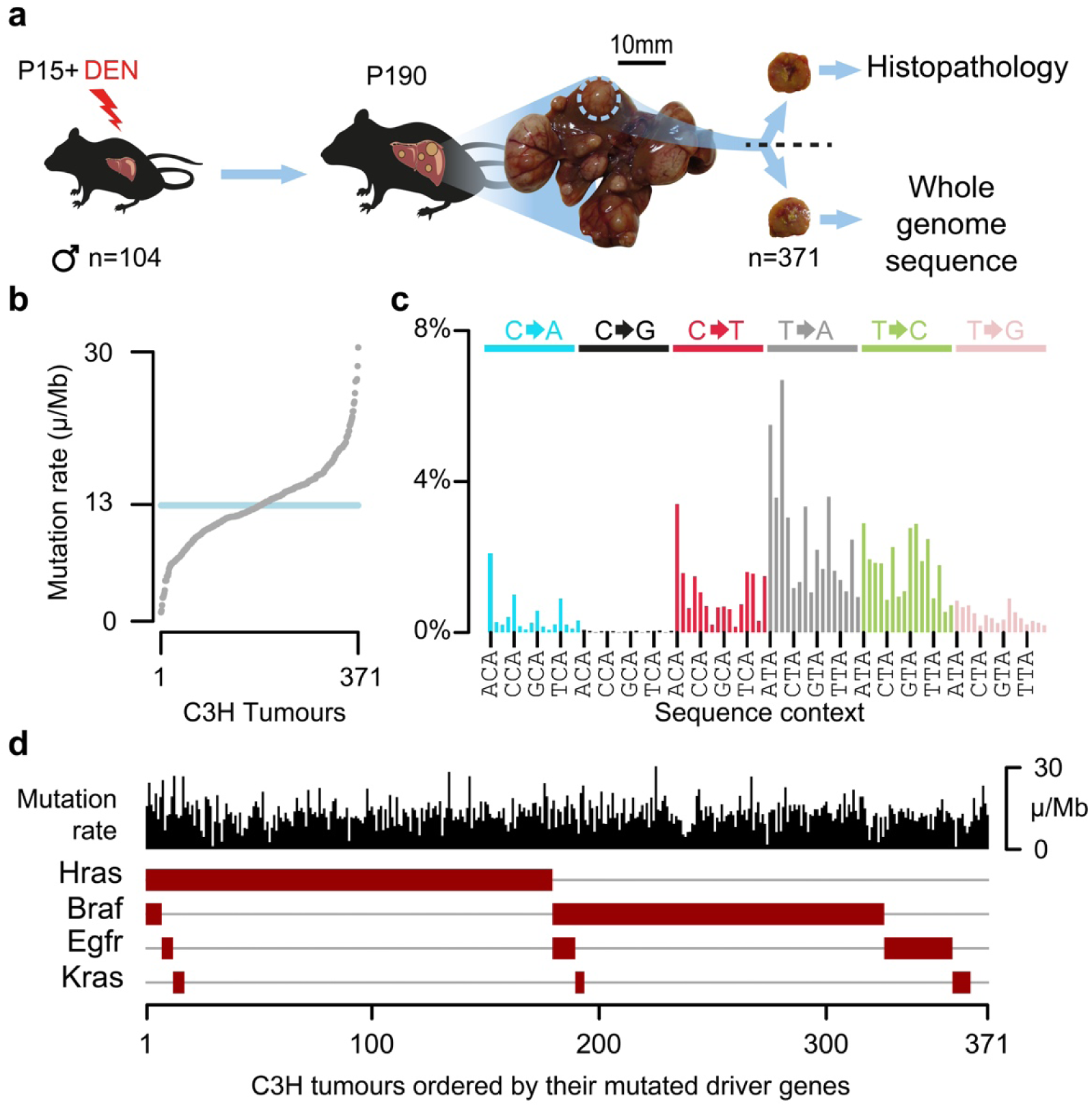
DEN-initiated tumours have a high burden of point mutations with a distinct mutation signature and driver mutations in the EGFR/RAS/RAF pathway. **a**, Fifteen-day-old (P15) male C3H/HeOuJ mice (n=104) received a single dose of diethylnitrosamine (DEN); tumours (n=371) were isolated 25 weeks after DEN treatment (P190), histologically analysed and subjected to whole genome sequencing. **b**, DEN-induced tumours displayed a median mutation rate of 13 mutations per million base pairs (μ/Mb). **c**, Mutation spectra histogram for the aggregated mutations of 371 C3H tumours showing the distribution of nucleotide substitutions, stratified by flanking nucleotide sequence context (96 categories). Sequence context for every fourth trinucleotide context is annotated (x-axis). **d**, Oncoplot summarising each tumour as a column with its mutation rate (black) and the presence of driver mutations in known driver genes (brown boxes). Tumours are ordered by the driver mutations identified.

Whole genome sequencing (WGS) of 371 independently-evolved tumours from 104 individual C3H mice (**Supplementary Table 1**) revealed that each diploid genome harboured ∼60,000 somatic point mutations (**Fig. 1b**), which equates to 13 mutations per megabase and is comparable to human cancers caused by exogenous mutagen exposure such as tobacco smoking and UV exposure^12, 13^. Insertion-deletion mutations, larger segmental changes, and aneuploidies were rare (**Extended Data Fig. 1a-f**). The tumour genomes were dominated (76%) by T→N/A→N mutations (where N represents any alternate nucleotide, **Fig. 1c**), consistent with previous studies implicating the long-lived thymine adduct O^4^-ethyl-deoxythymine as one of the principal mutagenic lesions generated by bioactivation of DEN in the liver by cytochrome P450 (CYP2E1)^14^. Deconvolution of mutation signatures revealed, in addition to the predominantly T→N signature (subsequently DEN1), a second signature (DEN2) that is typically present at a low level, but prominent in a minority of tumours (**Extended Data Fig. 1g-j**). DEN2 is characterised by C→T/G→A substitutions which likely represent the other principal mutagenic adduct of DEN^14^, O^6^-ethyl-2-deoxyguanosine, which can be repaired by the enzyme MGMT^15^.

Known driver mutations were identified in the EGFR/RAS/RAF pathway^10, 16, 17^ (**Fig. 1d**) validating this well controlled model system of liver tumourigenesis. These exhibited a strong propensity to be mutually exclusive: in 82% of C3H tumours only a single known driver mutation was observed. Similar results were replicated in CAST mice, though the proportion of driver gene usage differed (**Extended Data Fig. 1i,j**).

### Chromosome-scale segregation of lesions

Strikingly, in each tumour genome we observed multi-megabase segments with pronounced Watson versus Crick strand asymmetries of mutation spectra (**Fig. 2**). We define Watson strand bias as an excess of T→N over A→N mutations, when called on the forward strand of the reference genome, and the opposite as Crick strand bias. These biased regions show 23-fold (median) excess of their preferred mutation over its reverse complement. The asymmetric segments have a median span of 55Mb and often encompass an entire chromosome (**Fig. 2a-d**). The scale of these segmental asymmetries is orders of magnitude greater than those generated by TCR^9^, APOBEC mutagenesis^18, 19^ or produced by replication strand asymmetries^9, 20^. Despite the segmental strand asymmetry, the mutation load remains approximately uniform across the genome (**Fig. 2e**) and asymmetric segments do not correspond to deletions or other changes in DNA copy-number (**Fig. 2f**).

**Fig. 2.**
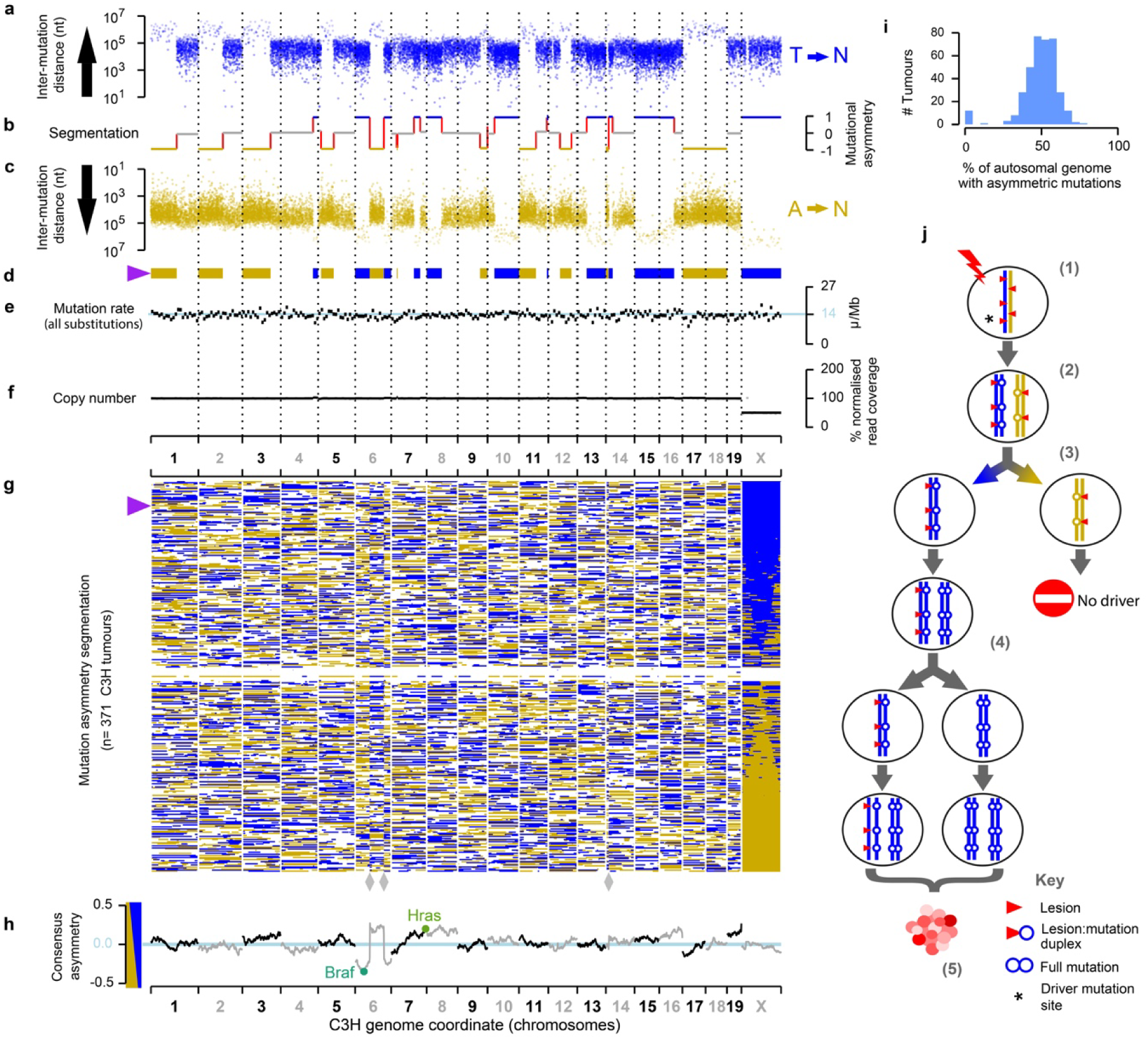
Chromosome-scale and strand asymmetric segregation of DNA lesions. **a-f**, An example DEN-induced C3H tumour (identifier: 94315_N8) with the genome shown over the x-axis. **a**-**c**, Mutational asymmetry. Individual T→N mutations shown as points, blue (T on the Watson strand, **a**) and gold (T on the Crick strand, **c**), the y-axis representing the distance to the nearest neighboring T→N mutation on the same strand. **b**, Segmentation of mutation strand asymmetry patterns. Y-axis position shows the degree of asymmetry (no bias: grey), and mutational symmetry switches indicated as red lines. **d**, Segmentation profile summarised as ribbon showing only the asymmetric segments. **e**, Mutation rate in 10Mb windows, blue line shows the genome wide rate for this tumour. **f**, DNA copy number in 10Mb windows (grey) and for each asymmetry segment (black). **g**, Summary ribbon plots (as in **d**) for all 371 C3H tumours, ranked by chromosome X asymmetry. Purple triangle indicates tumour shown in panels **a-f**. Reference genome mis-assembly points marked (grey diamonds). **h**, Balance of Watson versus Crick asymmetry amongst tumours, showing deviations at driver genes (calculation in 10Mb windows). **i**, Tumours consistently show segmental mutational asymmetry across 50% of their autosomal genome. **j**, Model for DNA lesion segregation as a mechanism to generate mutational asymmetries. The exposure of a mutagen generates lesions (red triangles) on both strands of the DNA duplex *(1)*. If not removed before or during replication *(2)* those lesions will segregate into two sister chromatids, one (blue) carrying only Watson strand lesions and subsequent templated errors, and the second (gold) only Crick strand lesions and their induced errors. Following mitosis, the daughter cells will have a non-overlapping complement of mutagen-induced lesions and resulting replication errors *(3)*, which are resolved into full mutations in the next round of replication *(4)*. The lesion containing strands segregate, becoming a progressively diminishing fraction of the lineage, yet continue as a template for replication. Only cell lineages containing cancer driver changes (* in *step* (*1)*) will expand into substantial clonal populations *(5)*.

This pervasive, strand-asymmetric mutagenesis can be explained as the consequence of DEN-induced lesions remaining unrepaired prior to genome replication. The first round of genome replication after DEN exposure results in two sister chromatids with independent lesions on their parent strands (**Fig. 2j**). The daughter strand is produced using a lesion-containing template whose complement is synthesised with reduced replication fidelity over damaged nucleotides, resulting in nucleotide misincorporation errors complementary to the lesion sites. These two sister chromatids necessarily segregate into separate daughter cells during mitosis and the heteroduplexes of lesions with paired mismatches are resolved into full mutations by subsequent replication cycles (**Fig. 2j**). We subsequently refer to this phenomenon as “lesion segregation”.

The haploid X chromosome always contains segments with either a strong Watson or Crick bias. On autosomal chromosomes, in addition to Watson or Crick biases, we also observe an unbiased state (**Fig. 2g**). We interpret this unbiased state as the aggregated biases of the two allelic autosomal chromosomes with conflicting strand asymmetries. More explicitly, when both copies of a chromosome have Watson strand bias, the genome shows a Watson strand bias (e.g. chromosome 15 in **Fig. 2a-c**); when one copy of the autosome has Watson bias and the other a Crick bias the two will cancel each other out and therefore appear unbiased (e.g. chromosome 19 in **Fig. 2a-c**). Under lesion segregation, these asymmetries represent the random retention of Watson or Crick strand biased segments over the whole genome, and are essentially the output of two independent 1:1 Bernoulli processes, analogous to two fair coin flips. In such a model, we would expect (1) 50% of the autosomal genome and (2) 100% of the haploid X chromosome to show mutational asymmetry; both predictions are supported by the observed data (**Fig. 2g,i**). A small fraction of tumours (3.5%) are outliers (**Fig. 2g,i**) with absent or muted mutational asymmetry; these features are associated with atypically low variant allele frequency distributions, indicating they may be polyclonal or polyploid (**Supplementary Table 1**).

The lesion segregation model predicts mutational asymmetries should span whole chromosomes, yet within a chromosome we commonly observe discrete switches between multi-megabase segments of Watson and Crick strand bias (**Fig. 2a-d,g**). Such switches likely represent sister chromatid exchanges resulting from homologous recombination (HR) mediated DNA repair events^21^ (**Extended Data Fig. 4a**) that are typically invisible to sequencing technologies because HR between sister chromatids is generally thought to be error-free^22^.

### HR repair increases genetic diversity

The observed rate of sister-chromatid exchange positively correlates with the genome-wide load of point mutations (**Extended Data Fig. 2a,b**). The presence of typically 27 (median) sister chromatid exchanges in each of 371 diploid tumour genomes meant we were well powered to detect recurrent exchange sites and biases in genomic context (**Extended Data Fig. 2c,d**). After filtering out three genomic locations that correspond to reference genome mis-assemblies (**Fig. 2g**; **Extended Data Fig. 3a,b**), we find that sister chromatid exchanges occur throughout the genome, with modest enrichment in transcriptionally inactive, late replicating regions and depletion in transcriptionally active and early replicating loci (**Extended Data Fig. 4b**).

**Fig. 3.**
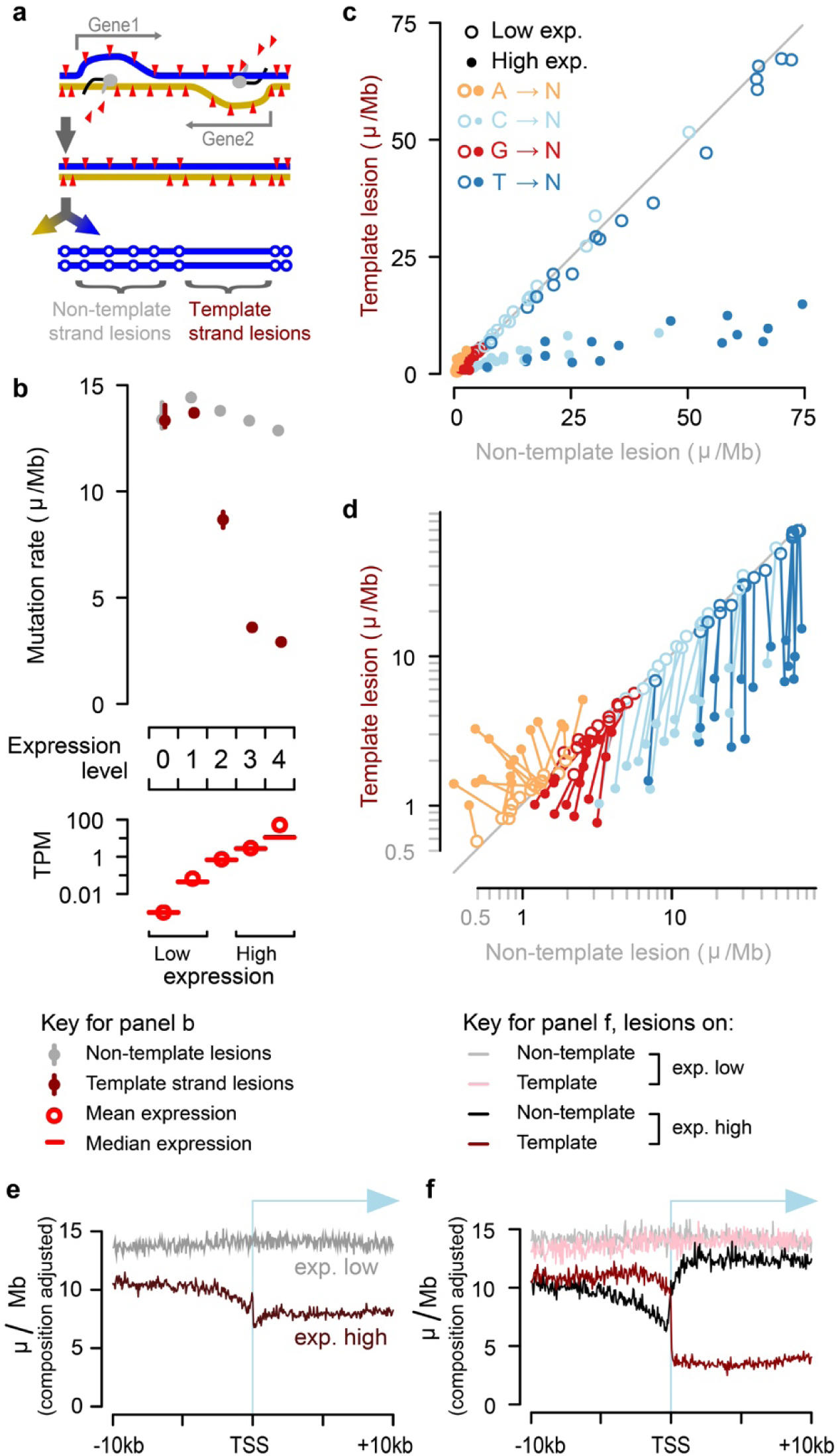
Identification of the lesion containing DNA strand allows processes such as transcription coupled repair (TCR) to be quantified with strand specificity. **a**, Transcription coupled repair of DNA lesions is expected to reduce the mutation rate only when lesions are on the template strand of an expressed gene. **b**, Transcription coupled repair of template strand lesions is dependent on transcription level (P15 liver, transcripts per million (TPM)). Confidence intervals (99%) are shown as whiskers, where broad enough to be visible. **c**, Comparison of mutation rates for the 64 trinucleotide contexts: each context has one point for low and one point for high expression. **d**, Data as in panel **c** plotted on log scale; there is a line linking low and high expression for the same trinucleotide context. **e**, Sequence composition normalised profiles of mutation rate around transcription start sites (TSS). **f**, Stratifying the data plotted in e by lesion strand reveals much greater detail, including the pronounced influence of bidirectional transcription initiation on the observed mutation patterns.

**Fig. 4.**
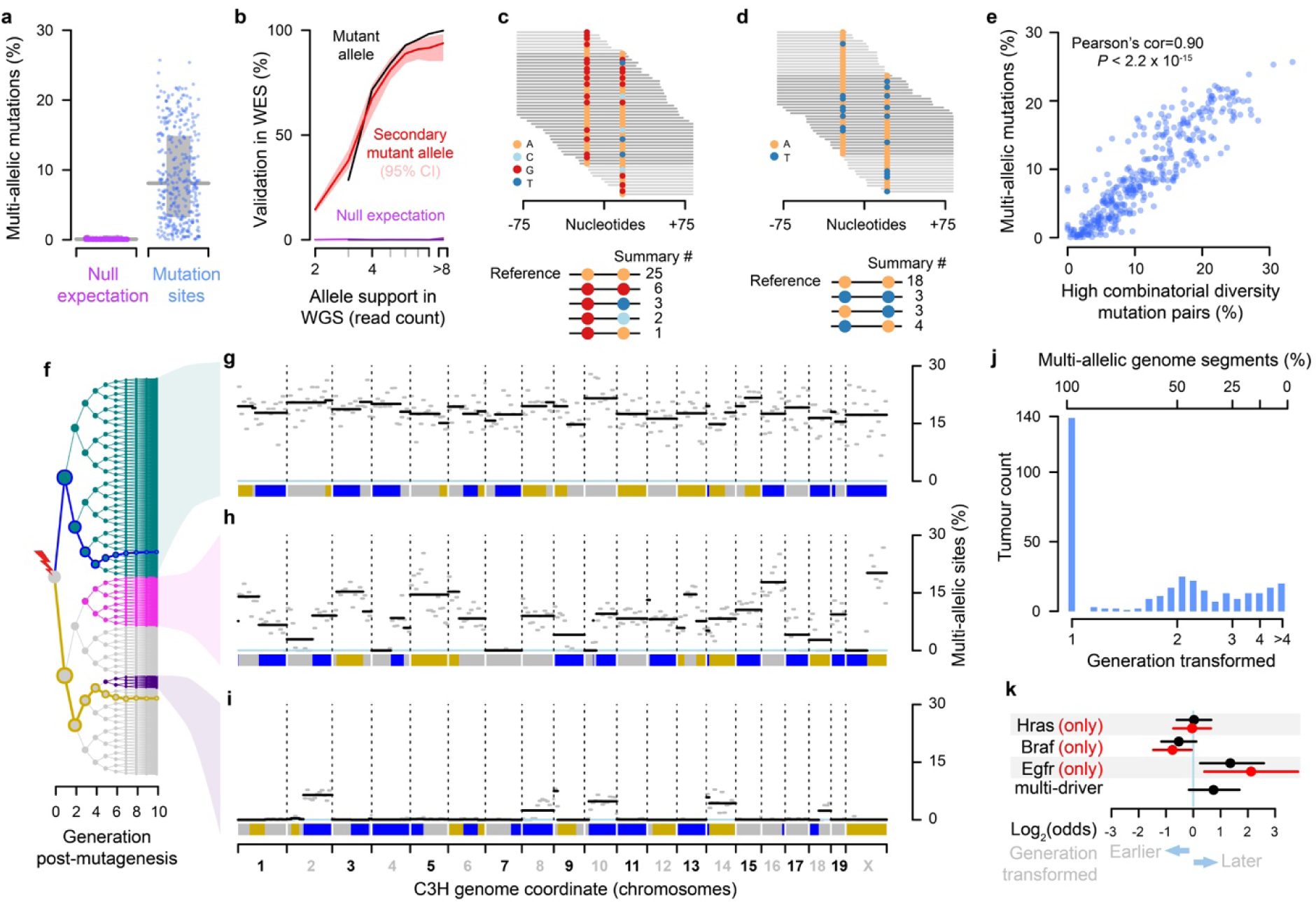
Lesion segregation generates multi-allelic and combinatorial genetic diversity. **a**, Percent of mutation sites with robust support for multi-allelic variation, one point per tumour. Grey line indicates median, and box the interquartile ranges. Null expectation (magenta) from permutation between tumours. **b**, Validation rate for whole genome sequence (WGS) mutation calls in replication whole exome sequencing (WES). Null expectation from permuting tumour identity between WGS and WES. **c**, Sequence reads over a pair of proximal mutations showing nucleotide calls per read; below summary count of read combinations. **d**, As for c showing combinatorial diversity between pairs of biallelic sites. **e**, Correlation between per-tumour multi-allelic rate and high combinatorial diversity mutation pairs (as in c,d), one point per tumour. **f**, Tree showing all possible progeny of a DEN mutagenised cell for the subsequent 10 generations. Blue and gold lines trace the simulated segregation of lesion-containing strands from the single-copy X chromosome. Coloured nodes show hypothetical transformation events and their daughter lineages that would give rise to the multi-allelic patterns in tumours shown to the right. **g-i**, Mutation asymmetry summary ribbons for example C3H tumours that show high **g**, variable **h**, or uniformly low **i** rates of genetic diversity; genome on the x-axis. The percent of mutation sites with robust support for multi-allelic variation calculated in 10Mb windows (grey) and for each asymmetric segment (black). **j**, Histogram of the estimated cell generation post-DEN exposure from which C3H tumours developed. **k**, Enrichment of specific driver gene mutations in earlier (generation 1) and later (generation >1) developing tumours. All tumours containing the indicated driver mutation (black); the subset of tumours with just the indicated driver and no other driver mutation (red) Multi-driver denotes all tumours that contain multiple identified driver genes in the EGFR/RAS/RAF pathway.

The fine mapping (∼20kb resolution) of sister chromatid exchange events allowed us to test the fidelity of HR between sister chromatids. Aggregate analysis indicated that sister chromatid exchange sites have approximately double the rate of point mutations compared to the remainder of the genome, suggesting HR may be an error-prone process. However the spectrum of point mutations in these regions did not differ from the rest of the tumour genomes (**Extended Data Fig. 4c-f**). A shift in spectrum would be expected if additional mutations had been introduced during the homologous repair process, independent of the chemically induced lesion repair. We propose that a model of Holliday intermediate branch migration can explain these observations (**Extended Data Fig. 4g** and would mean that, to the limit of our resolution, the HR process is error-free. Under this model the observed increase in mutation rate is actually a HR driven increase in mutational diversity within the cell population.

### Lesion segregation reveals oncogenic selection

The random segregation of sister chromatids into daughter cells would result in 50% Watson strand and 50% Crick strand lesion retention on average across tumours. The majority of the genome conforms to this prediction (**Fig. 2h**). We observe striking deviations from this 50:50 ratio at loci spanning known murine hepatocellular carcinoma driver genes (**Fig. 2h**). For example the *Braf* T→A mutation at codon 584 is a known oncogenic driver^10^ and observed in 153/371 of the C3H tumours. Presuming that the *Braf* mutation was DEN induced, we would expect the mutation to have occurred in a chromosomal segment that retained T-lesions on the same strand as the driver T→A change. Indeed this is the case (94%; 144/153 tumours retain lesions on the expected strand, Fisher’s exact test p=3.6×10^−19^, rejecting the 50:50 null expectation). In contrast, we would not expect tumours lacking the *Braf* mutation to show a systematic bias in the lesion strands retained at this genomic locus, and our observations again match expectations (47% Crick bias, 53% Watson bias, p=0.88, not rejecting the 50:50 null expectation). We applied this general test for oncogenic selection at sites with sufficient recurrent mutations to have statistical power. Our results confirmed significant oncogenic selection of previously identified driver mutations in *Hras*, *Braf* and *Egfr*, but did not support it at two other recurrently (n>33) mutated sites, demonstrating the specificity of this test (**Extended Data Table 1**).

### DNA repair with lesion strand resolution

Resolving DNA lesions to specific strands in a single mutagenised cell cycle presents a unique opportunity to investigate strand-specific interactions with DNA damage and repair *in vivo*, such as quantifying the efficiency and biases of TCR. Transcription-coupled nucleotide excision repair specifically removes DNA lesions from the mRNA template strand rather than from the non-template strand in expressed genes (**Fig. 3a**)^23, 24^. To explore this, we generated total transcriptomes of liver tissue from P15 C3H and CAST mice, corresponding to the known tissue of origin as well as the exact developmental timing of DEN mutagenesis.

For each gene in each tumour, we resolved whether the lesion strand was the mRNA template or non-template strand, and calculated mutation rates stratified by both expression level and lesion strand (**Fig. 3b**). As expected, TCR was highly specific to the template strand and correlated closely with gene expression. Among genes without detectable expression, there was no reduction or observable transcription strand-bias in the mutation rate. In contrast, the mutation rate of the most highly expressed genes was reduced by 79.8±1.0% if the tumour inherited lesions on the template strand. We also detect a small transcription-associated decrease in mutation rate for lesions on the non-template strand: 10.7±1.4% reduction in rate relative to lowly-expressed genes.

We next considered the specificity of TCR, comparing the rates of mutation for each trinucleotide context (n=64 categories) between template and non-template strands, stratified by expression level (**Fig. 3c,d**). All measures are rates, thus taking into account sequence composition differences between sets of genes and DNA strands. The most common mutations (T→N), have an 82% (s.d. 6.8% across sequence contexts) lower rate on the template strand than the non-template strand for highly expressed genes; the non-template mutation rate is the same regardless of expression level (**Fig. 3d**, dark-blue lines are close to vertical), consistent with expectations^23^.

Mutations from C and G on the template strand also show a high efficiency of TCR (70% (s.d. 7.8%) and 34% (s.d. 21%) respectively, **Fig. 3d**). For these mutation classes, however, there is also a consistent transcription-dependent reduction of mutation rate when the lesions are inferred to be on the non-template strand (lines are deflected left, rather than vertical). This may indicate the targeting of non-transcription coupled repair processes to accessible genic regions. Notably, though comparatively rare, mutations from adenine on the lesion containing strand are increased with transcription (**Fig. 3d**). This unexpected observation could be due to the activity of error-prone trans-lesion DNA polymerase Pol-η which targets transcribed regions, where it is known to promote mutations specifically at A:T base-pairs^25^.

Prior analyses of transcription coupled repair could not resolve the lesion containing strand^9, 23, 26^. Consistent with these previous findings, we observe reduced mutation rates broadly across the transcription start site (TSS) region and into the body of active genes (**Fig. 3e**). A notable feature of this profile is the relative increase in mutation rate for the core promoter located in the 200 nucleotides immediately upstream of the TSS^27^. Including lesion strand information in the analysis (**Fig. 3f**) shows the relative increase in mutation rate over the core promoter to be a result of high rates of TCR upstream and downstream, but a relative depletion of TCR activity over the promoter itself, results that are replicated in CAST mice (**Extended Data Fig. 5a-e**). The ability to resolve the lesion strand in our analysis (**Fig. 3f**) newly reveals the striking and distinct contributions of bidirectional transcription from active promoters^28^ in shaping the observed mutation patterns.

**Fig. 5.**
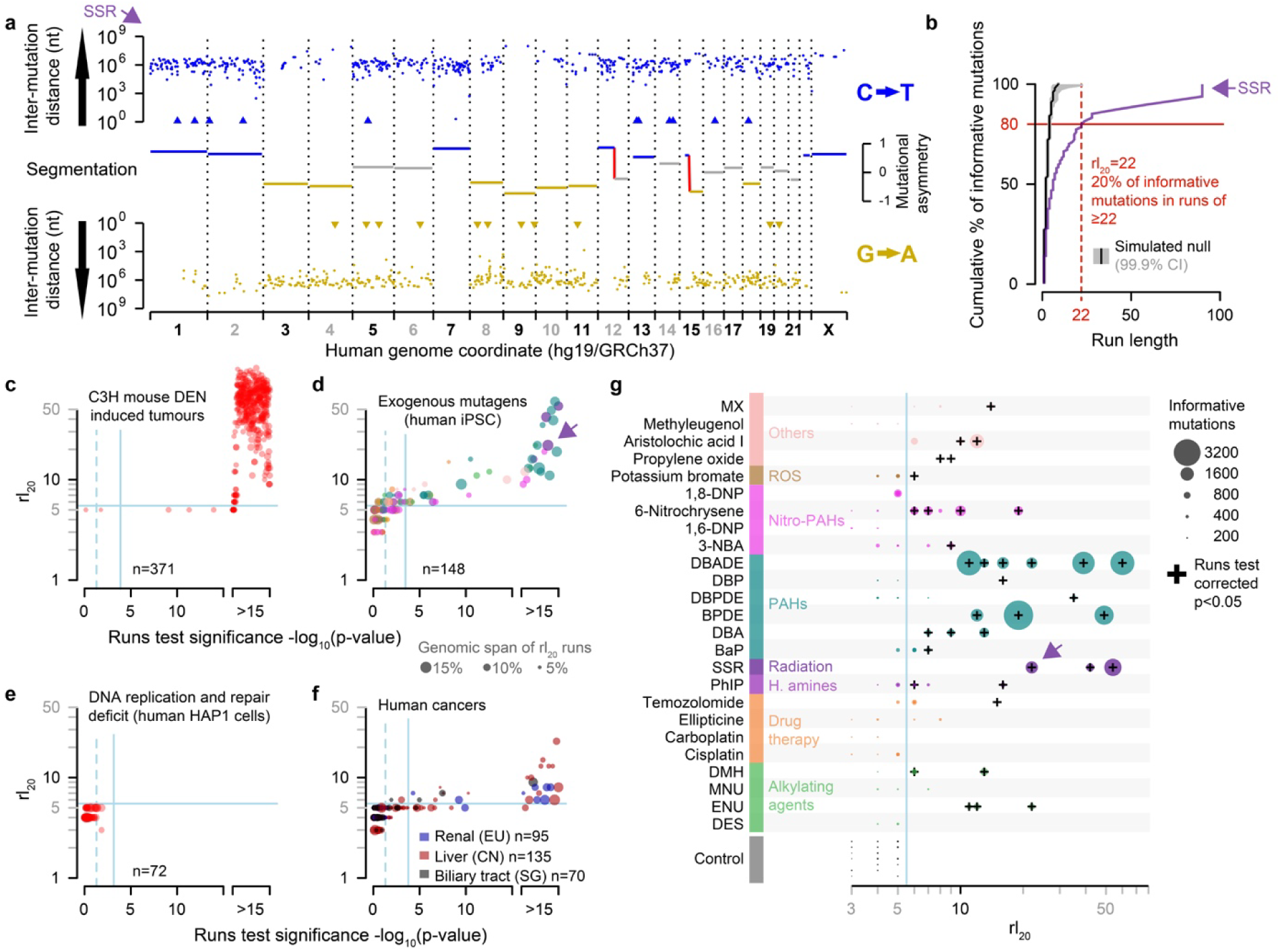
Lesion segregation is a pervasive feature of exogenous mutagens and is evident in human cancers. **a**, Mutational asymmetry in human cells subject to simulated solar radiation (SSR)^5^, highlighted subsequently with a purple arrow. Dinucleotide changes (CC→TT, GG→AA) are shown as triangles. **b**, The runs-based rl_20_ metric, calculated for the clone shown in panel **a**; here, 20% of informative mutations (C→T/G→A) are in strand asymmetric runs of 22 consecutive mutations or longer (e.g. ≥22 C→T without an intervening G→A). **c-f**, The rl_20_ metric and runs tests. Solid blue lines show Bonferroni adjusted p=0.05 thresholds, p-values < 1×10^−15^ are rank-ordered. **c**, DEN-induced C3H tumours. **d**, Mutagen exposed human cells^5^, colour corresponds to the mutagen key in panel **g**. **e**, Cell-lines with genetically perturbed genome replication and maintenance machinery^30^. **f**, Human cancers from International Cancer Genome Consortium projects. **g**, All 25 mutagens identified as producing robust mutation spectra when human induced pluripotent stem cells are exposed^5^, grouped by type of agent. See **Supplementary Table 2** for the full details of abbreviated mutagen exposures. The rl_20_ metric (x-axis) is plotted for each replicate clone, the size of each data point is scaled to the number of informative mutations in the clone.

### Lesion segregation generates genetic diversity

The lesion segregation model (**Fig. 2j**) predicts that a segregating lesion may template multiple rounds of replication in successive cell cycles. In such a scenario, each replication across the lesion could incorporate different incorrectly paired nucleotides – or even the correctly paired nucleotide – opposite a persistent lesion. Consistent with this notion, hundreds of multi-allelic mutations have recently been reported from single cell sequencing of human cancer samples^29^.

We investigated the extent of multi-allelic variation within our data by analysing the sequencing reads at the sites of identified mutations. For example, a nucleotide position with multiple high-confidence reads supporting a reference T, mutation to A and also mutation to C would be considered multi-allelic. We find that on average 8% of mutated sites in DEN induced tumours exhibit evidence of multi-allelic variants (n=1.8 million sites across our C3H data set), though this value ranges from <1% to 25.7% between tumours (**Fig. 4a**). As a control we performed equivalent analysis on sites that had been called as mutated in a randomly selected proxy tumour (**Methods**). On average, only 0.098% (95% CI: 0.043-0.25%) of these control sites show evidence of non-reference nucleotides by the same criteria.

We further validated the multi-allelic variant calls from whole genome sequencing within independently performed deep exome sequencing of the same tumours^10^. The second and subsequent alternate alleles show the same profile of read depth-dependent validation rate as the called mutant allele, and clear separation from control analyses with mis-paired exome and genome sequence (**Fig. 4b**).

The independent generation of multi-allelic variation at sites across the genome produces combinatorial genetic diversity that is not expected under a model of purely clonal expansion. This combinatorial diversity can be directly visualised in pairs of mutated sites close enough to be spanned by individual sequencing reads (**Fig. 4c,d**). Analysis of such reads demonstrates the presence of combinations of alleles between bi-allelic sites that requires lesions to have been replicated over, without generating a mutation in some cell divisions (**Fig. 4d**). As expected for orthogonal measures of the generated genetic diversity, the tumour-wide level of combinatorial diversity from such proximal mutation pairs is closely correlated with the multi-allelic rate (**Fig. 4e**), and highlights the consistently high variance of these measures between tumours.

The explanation for this inter-tumour variance becomes evident when plotting the distribution of multi-allelism along each tumour genome (**Fig. 4f-i**). Tumours with high rates of genetic diversity typically have uniformly high rates of multi-allelism across their genome (**Fig. 4g**). They likely developed from a first generation daughter of the original DEN mutagenised cell, in which all DNA is a duplex of a lesion containing and non-lesion containing strand. Replication over lesion containing strands in subsequent generations therefore produces multi-allelic variation at a uniform rate throughout the genome.

Tumours with lower total levels of genetic diversity exhibit discrete genomic segments of high and low multi-allelism (**Fig. 4h,i**). These tumours can be explained as having developed from a cell a few generations subsequent to DEN treatment. Each mitosis following DEN exposure is expected to dilute the lesion containing strands present in each daughter cell by approximately 50% per generation, assuming random segregation. Only fractions of the genome retaining the lesions will go on to generate multi-allelic and combinatorial genetic diversity in the daughter lineages. As expected from the lesion segregation model with sister chromatid exchange, the multi-allelic patterns mirror the mutational asymmetry segmentation pattern. Tumours with uniformly low levels of multi-allelism can similarly be explained as having developed from a still later generation cell that had not inherited any lesion containing strands.

By estimating the fraction of chromosomal segments that are multi-allelic, we can infer the cell generation post-DEN exposure that the tumour grew from (**Fig. 4j**). This indicates that 67% of C3H and 21% of CAST tumours developed from the first generation daughter cell following DEN exposure, indicating the single large burst of mutations was instantly transformative. For the remainder, the observed fractions of multi-allelic segments cluster around expectations for second and subsequent cell generations, suggesting that either an additional mutational hit, the generation of a specific mutation allele combination, or an external trigger is required for transformation. Intriguingly, there is a bias in driver gene usage between early and late transforming tumours (**Fig. 4k**) with *Egfr* driven tumours significantly later to transform. Analysis of the mutational landscape from lesion segregation revealed novel mechanisms of generating and sampling genetic diversity, as well as giving insights into the earliest events in oncogenesis.

### Lesion segregation is ubiquitous

Lesion segregation is a major feature of DEN mutagenesis in mouse liver. This immediately raises two important questions: are other DNA damaging agents also characterised by lesion segregation? And does lesion segregation occur in human cells and cancers? A recent study exposed human induced pluripotent stem cells (iPSCs) to 79 known or suspected environmental mutagens and found 41 of them produced nucleotide substitutions above background expectations^5^. Although not previously noted in these data, we found that many of these mutagenic exposures generated chromosome-scale lesion segregation patterns in the same manner as our *in vivo* DEN model (**Fig. 5a**; **Extended Data Fig. 6a-d**).

To allow generalised searching of genomic mutation data, we identified the most common nucleotide substitution type in each sample (e.g. C→T/G→A in simulated solar radiation exposure, **Fig. 5a**) and applied runs-based tests to quantify the segmental asymmetry of those informative mutations. The application of runs-based tests (e.g. the rl_20_ metric) (**Fig. 5b-f**) revealed that segmental mutational asymmetry is a common feature of DNA damaging mutagens (**Fig. 5d**). We detect significant mutational asymmetry in every sample with good statistical power (>1,000 informative mutations, **Fig. 5g**), including multiple clinically relevant insults, including sunlight (simulated solar radiation, SSR), tobacco smoke (BPDE) and chemotherapeutic agents (temozolomide). We conclude that the chromosome-scale segregation of lesions and the resulting strand asymmetry of mutation patterns is a general feature of all tested DNA damaging mutagens.

In prior work from the same laboratory as Kucab *et al*., components of the mismatch repair pathway were disabled and other mutator phenotypes were genetically engineered. These experiments all generate mutations through mechanisms that are independent of DNA damage^30^. Cells were grown from single clones, sequenced and analysed as for the iPSC mutagenesis data, and showed a similar range of mutation counts. As their mutations are independent of DNA lesions, they were not expected to show lesion segregation patterns. Confirming this expectation no significant asymmetry was detected (**Fig. 5e**).

A common feature of our DEN mutagenesis experiment and mutagen exposure in human IPS cells^5^ is the application of a single mutagenic insult. Our lesion segregation model predicts that mutagen exposures over multiple cell cycles would progressively mask the mutational strand asymmetry pattern as independent patterns are successively superimposed. Most exogenous mutagens implicated in the pathogenesis of human cancers are encountered as repeated episodic exposures, for example smoking or UV exposure. Therefore, even though we have shown that UV exposure does cause lesion segregation in human cells (**Fig. 5a**) it is unlikely that the mutational asymmetry diagnostic of lesion segregation would be detected in skin cancers.

Despite the low prior expectation of detecting lesion segregation patterns in clinical human cancers, we used the same algorithm as for human iPS cells to analyse the cancer genomes from the International Cancer Genome Consortium^3^ (n=19,778 cancers from 22 cancer primary sites). This analysis identified multiple cancers that are highly significant (**Fig. 5f**) and clearly show mutational asymmetry patterns characteristic of lesion segregation (**Extended Data Fig. 7a-c**). The majority of these tumours are renal, hepatic or biliary in origin, and show a high mutation rate and strand asymmetry of T→A/A→T mutations, consistent with known aristolochic acid exposure^3^ (**Supplementary Table 2**). We conclude that lesion segregation is likely to be a general feature of mammalian cells with episodic or continuous exposure to mutagens, though visualised most clearly in homogenous cell populations subjected to single doses of a mutagen.

## Discussion

Here we show that most mutagenic DNA lesions are not resolved as mutations within a single cell-cycle. Instead, lesions segregate unrepaired into daughter cells for multiple cellular generations, resulting in the chromosome-scale strand asymmetry of subsequent mutations. Initially discovered in a well-powered *in vivo* model of oncogenesis, we also demonstrate that lesion segregation is ubiquitous to all tested mutagens, occurs in human cells, and is evident in human cancers. The striking pattern of mutation asymmetry that we observe occurs as a result of a single mutagenic treatment. Since most cancers are subject to multiple damaging events over their life history, they will acquire new waves of segregating lesions after each exposure, thus resulting in mutually confounding mutation patterns. Hence, although the mutation asymmetry pattern is most pronounced in bulk sequencing of liver and kidney cancers in human cohorts, lesion segregation has likely shaped all genomes that have suffered DNA damage, which has important implications for tumour evolution and heterogeneity.

The lesion segregation model makes several predictions, which we confirmed in our experimental results in both C3H and CAST mice, and are further validated in published human datasets. As predicted for the random segregation of lesion strands, segmental asymmetry is present throughout the haploid X chromosome and 50% of the remaining genome. While the genome-wide pattern of strand retention within and between tumours is random, it is focally biased over regions containing driver genes, which itself provides a novel approach to detect oncogenic selection. The persistence of lesions for more than one cellular generation predicts the presence of multi-allelic sites, which are abundant in our *in vivo* experimental system, and were recently described in smaller numbers in human cancers^29^ and a well-controlled cell lineage tracking system^31^.

Our discovery of pervasive lesion segregation challenges long standing assumptions in the analysis of cancer evolution^32^. For example, the widely used infinite sites model^33^ does not allow for recurrent rounds of mutation at the same site, such as those generated by segregating lesions. It also provides new opportunities for understanding cancer evolution, through the use of the mutational asymmetry and multi-allelic-rate patterns to track events during oncogenesis and, in some instances, to quantify selection. A far-reaching implication of lesion segregation is that it may provide a window of opportunity for a cancer to sample the fitness of mutation combinations within the lineage, circumventing Muller’s ratchet^34^ and Hill-Robertson interference: low efficiency of selection due to the inability to separate mutations of opposing fitness effects^35, 36^. Consequently, DNA damaging chemotherapeutics, particularly large or closely spaced doses that generate persistent lesions, could inadvertently provide an opportunity for efficient selection of the resulting mutations. This insight may guide the development of more effective chemotherapeutic regimens.

Consistent with our initial motivation, the ability to resolve lesion strandedness has enabled more refined analyses of DNA damage and repair processes. For example, our analysis of transcription coupled repair revealed with unforeseen clarity how frequent bidirectional transcription shapes the distribution of mutations at transcription start sites. Our data also afforded high resolution mapping of sister chromatid exchange events, quantifying for the first time the increased mutational diversity at sites of homology driven repair. Both of these practical applications of lesion segregation provide new vistas for the exploration of genome maintenance and fundamental molecular biology.

Once identified, lesion segregation is a deeply intuitive concept. DNA damage and lesion formation occurs independently on the Watson and Crick strands and those surviving DNA repair persist through mitosis. Replication over these segregating lesions generates combinatorial genetic diversity, thus providing opportunities for selection and adaptation long assumed to be impossible in a clonally expanding population. The discovery of pervasive lesion segregation profoundly revises our understanding of how the architecture of DNA repair and clonal proliferation can conspire to shape the cancer genome.

## Methods

### Mouse colony management

Animal experimentation was carried out in accordance with the Animals (Scientific Procedures) Act 1986 (United Kingdom) and with the approval of the Cancer Research UK Cambridge Institute Animal Welfare and Ethical Review Body (AWERB). Animals were maintained using standard husbandry: mice were group housed in Tecniplast GM500 IVC cages with a 12-hour light / 12-hour dark cycle and *ad libitum* access to water, food (LabDiet 5058), and enrichments.

### Chemical model of hepatocarcinogenesis

15-day-old (P15) male C3H and CAST mice were treated with a single intraperitoneal (IP) injection of N-Nitrosodiethylamine (DEN; Sigma-Aldrich N0258; 20 mg/kg body weight) diluted in 0.85% saline. Liver tumour samples were collected from DEN-treated mice 25 weeks (C3H) or 38 weeks (CAST) after treatment. All macroscopically identified tumours were isolated and processed in parallel for DNA extraction and histopathological examination. Non-tumour tissue from untreated P15 mice (ear, tail, and background liver) was sampled for control experiments.

### Tissue collection and processing

Liver tumours of sufficient size (≥2 mm diameter) were bisected; one half was flash frozen in liquid nitrogen and stored at −80°C for DNA extraction, and the other half was processed for histology. Tissue samples for histology were fixed in 10% neutral buffered formalin for 24 h, transferred to 70% ethanol, machine processed (Leica ASP300 Tissue Processor; Leica, Wetzlar, Germany), and paraffin embedded. All formalin-fixed paraffin-embedded (FFPE) sections were 3 μm in thickness.

### Histochemical staining

FFPE tissue sections were haematoxylin and eosin (H&E) stained using standard laboratory techniques. Histochemical staining was performed using the automated Leica ST5020; mounting was performed on the Leica CV5030.

### Imaging

Tissue sections were digitised using the Aperio XT system (Leica Biosystems) at 20x resolution; all H&E images are available in the BioStudies archive at EMBL-EBI under accession S-BSST129.

### Tumour histopathology

H&E sections of liver tumours were blinded and assessed twice by a pathologist (S.J.A); discordant results were reviewed by an independent hepatobiliary pathologist (S.E.D). Tumours were classified according to the International Harmonization of Nomenclature and Diagnostic Criteria for Lesions in Rats and Mice (INHAND) guidelines ^37^. In addition, tumour grade, size, morphological subtype, nature of steatosis, and mitotic index were assessed (**Supplementary Table 1**), as well as the presence of cystic change, haemorrhage, necrosis, or vascular invasion.

### Sample selection for WGS

Tumours which met the following histological criteria were selected for whole genome sequencing (C3H n=371, CAST n=84): (i) diagnosis of either dysplastic nodule (DN) or hepatocellular carcinoma (HCC), (ii) homogenous tumour morphology, (iii) tumour cell percentage >70%, and (iv) adequate tissue for DNA extraction. Neoplasms with extensive necrosis, mixed tumour types, a nodule-in-nodule appearance (indicative of an HCC arising within a DN), or contamination by normal liver tissue were excluded. Since carcinogen-induced tumours arising in the same liver are independent ^10^, multiple tumours were selected from each mouse to minimise the number of animals used. A subset of normal (non-tumour) samples from untreated mice were also sequenced (C3H n=13, CAST n=7).

### Whole genome sequencing

Genomic DNA was isolated from liver tissue and liver tumours using the AllPrep 96 DNA/RNA Kit (Qiagen, 80311) according to the manufacturer’s instructions. DNA quality was assessed on a 1% agarose gel and quantified using the Quant-IT dsDNA Broad Range Kit (Thermo Fisher Scientific). Genomic DNA was sheared using a Covaris LE220 focused-ultrasonicator to a 450 bp mean insert size.

WGS libraies were generated from 1 μg of 50 ng/ul high molecular weight gDNA using the TruSeq PCR-free Library Prep Kit (Illumina), according to the manufacturer’s instructions. Library fragment size was determined using a Caliper GX Touch with a HT DNA 1k/12K/Hi Sensitivity LabChip and HT DNA Hi Sensitivity Reagent Kit to ensure 300-800 bp (target ∼450 bp).

Libraries were quantified by real-time PCR using the Kapa library quantification kit (Kapa Biosystems) on a Roche LightCycler 480. 0.75 nM libraries were pooled in 6-plex and sequenced on a HiSeq X Ten (Illumina) to produce paired-end 150 bp reads. Each pool of 6 libraries was sequenced over eight lanes (minimum of 40x coverage).

### Variant calling and somatic mutation filtering

Sequencing reads were aligned to respective genome assemblies (C3H = C3H_HeJ_v1; CAST = CAST_EiJ_v1) ^38^ with bwa-mem (v.0.7.12) ^39^ using default parameters. Reads were annotated to read groups using the picard (v.1.124) ^40^ tool AddOrReplaceReadGroups, and minor annotation inconsistencies corrected using the picard CleanSam and FixMateInformation tools .Bam files were merged as necessary, and duplicate reads were annotated using the picard tool MarkDuplicates.

Single nucleotide variants were called using Strelka2 (v.2.8.4) ^41^ implementing default parameters. Initial variant annotation was performed with the GATK (v.3.8.0) ^42^ walker CalculateSNVMetrics ^43^. Genotype calls with a variant allele frequency < 0.025 were removed. Although inbred strains were used, fixed genetic differences between the colonies and the reference genome, as well as small numbers of germline variants segregating within the colonies were identified. For each strain, fixed differences were identified as homozygous changes present in 100% of genotyped samples were filtered out. Segregating variants were filtered based on the excess clustering of mutations to animals with shared mothers. To generate a null expectation taking into account the family structure of the colonies, the parent-offspring relationships were randomly permuted 1,000 times. For each count of recurrent mutation (range 5 to 371 inclusive), we determined the null distribution of expected distinct mothers. Comparing this to the observed count of distinct mothers for each recurrent (n>4) mutation, those with a low probability (p<1×10^−4^, pnorm function from R (v.3.5.1) ^44^) under the null were excluded from analyses.

Copy number variation between tumours within strains was called using CNVkit (v.0.9.6) ^45^. Non-tumour reference coverage was provided from non-tumour control WGS data (C3H n=11, CAST n=7) and per tumour cellularity estimates (see below) were provided.

### RNA-sequencing

Total RNA was extracted from P15 liver tissue (n=4 biological replicates per strain) using QIAzol Lysis Reagent (Qiagen), according to the manufacturer’s instructions. DNase treatment and removal were performed using the TURBO DNA-freeTM Kit (Ambion, Life Technologies), according to the manufacturer’s instructions. RNA concentration was measured using a NanoDrop spectrophotometer (Thermo Fisher); RNA integrity was assessed on a Total RNA Nano Chip Bioanalyzer (Agilent)

Total RNA (1 μg) was used to generate sequencing libraries using the TruSeq Stranded Total RNA Library Prep Kit with Ribo-Zero Gold (Illumina), according to the manufacturer’s instructions. Library fragment size was determined using a 2100 Bioanalyzer (Agilent). Libraries were quantified by qPCR (Kapa Biosystems). Pooled libraries were sequenced on a HiSeq4000 to produce ≥40 million paired-end 150 bp reads per library.

### RNA-seq data processing and analysis

Transcript abundances were quantified with Kallisto (v.0.43.1)^46^ (using the flag --bias) and a transcriptome index compiled from coding and non-coding cDNA sequences defined in Ensembl v91^47^. Transcripts per million (TPM) estimates were generated for each annotated transcript and summed across alternate transcripts of the same gene for gene-level analysis. Transcription start sites (TSS) for each gene were annotated with Ensembl v91 and based upon the most abundantly expressed transcript. RNA-seq data are available at Array Express at EMBL-EBI under accession E-MTAB-8518.

### Genomic annotation data

Mouse liver proximity ligation sequencing (HiC) data were downloaded from GEO (GSE65126) ^48^, replicates were combined, then aligned to GRCm38 ^49^ and processed using the Juicebox (v.7.5) and Juicer scripts ^50^ to obtain the HiC matrix. Eigenvectors were obtained for 500kb consecutive genomic windows over each chromosome from the HiC matrix using Juicebox and subsequently oriented (to distinguish compartment A from B) using GC content per 500kb bin. We used progressiveCactus ^51^ to project the 500kb windows into the C3H reference genome and Bedtools (v.2.28.0) to merge syntenic loci between 450 and 550 kb in size, removing the second instance where we observed overlaps.

Genic annotation was obtained from Ensembl v91 ^47^for the corresponding C3H and CAST reference genome assemblies (C3H_HeJ_v1, CAST_EiJ_v1). Genomic repeat elements were annotated using RepeatMasker (v.20170127) ^52^ with the default parameters and libraries for mouse annotation.

### The analysable fraction of the genome

Analysis and sequence composition calculations were confined to the main chromosome assemblies of the reference genome (chromosomes 1-19 and X). Using WGS of non-tumour liver, ear and tail samples (C3H n=11, CAST n=7) collected and sequenced contemporaneously with tumour samples, genome sequencing coverage was calculated for 1kb windows using multicov in Bedtools (v.2.28.0) ^53^. Windows with read coverage >2 s.d. from the autosomal mean were flagged as suspect in each tumour. Read coverage over the X chromosome was doubled in these calculations to account for the expected hemizygosity in these male mice. Any 1kb window identified as suspect in >90% of these non-tumour samples was flagged as “abnormal read coverage” (ARC) and masked from subsequent analysis. This masked 12.7% of the C3H and 11.5% of the CAST reference genomes yielding analysable haploid genome sizes of C3H = 2,333,783,789 nt and CAST = 2,331,370,397 nt.

### Mutation rate calculations

Mutation rates were calculated as 192 category vectors representing every possible single nucleotide substitution conditioned on the identity of the upstream and downstream nucleotides. Each rate being the observed count of a mutation category divided by the count of the trinucleotide context in the analysed sequence. To report a single aggregate mutation rate, the three rates for each trinucleotide context were summed to give a 64 category vector and the weighted mean of that vector reported as the mutation rate. The vector of weights being the trinucleotide sequence frequency of a reference sequence, for example the composition of the whole genome. In the case of whole genome analysis, the same trinucleotide counts are used in (1) the individual category rates calculation and (2) the weighted mean of the rates, cancelling out. For windowed comparisons of mutation rates, the weighted mean is calculated using the genome wide composition of trinucleotides rather than the local sequence composition, providing a compositionally adjusted mutation rate estimate. For mutation rates in TCR analysis, the same compositional adjustment was carried out but using the trinucleotide composition of the aggregate genic spans of genome (minus ARC regions) for normalisation.

### Mutation signatures

The 96 category “folded” mutation counts for each of the 371 C3H tumours were deconvolved into the best fitting number (*K*) of component signatures using sigFit (v.2.0) ^54^ with 1,000 iterations and *K* set to integers 2 to 8 inclusive. A heuristic goodness of fit score based on cosine similarity favoured instances where *K*=2. The DEN1 and DEN2 signatures reported were obtained by running sigFit with 30,000 iterations for *K*=2. Analysis of CAST tumours gave less distinct separation of signatures so the C3H derived DEN1 and DEN2 were used for both strains. To fit signatures to each tumour we used sigFit provided with the DEN signatures and additional SPONT1 and SPONT2 signatures that were derived from equivalent WGS analysis of spontaneous (non-DEN induced) C3H tumours.

### Driver mutation identification

Driver mutations in the known oncogenic driver genes *Egfr*, *Braf*, *Hras*, and *Kras* were previously identified for C3H ^10^ and the orthologous mutations identified in CAST derived tumours were annotated as driver mutations.

### Mutational asymmetry segmentation and scoring

For each tumour a focal subset of “informative” mutation types were defined, T→N/A→N mutations, in the case of DEN-induced tumours. The order of focal mutations along each chromosome was represented as a binary vector (e.g. 0 for T→N, 1 for A→N). Vectors corresponding to each chromosome of each tumour were processed with the cpt.mean function of the R Changepoint (v.2.2.2) ^55^ package run with an Akaike information criterion (AIC) penalty function, maximum number of changepoints set to 12 (Q=12), and implementing the PELT algorithm for optimal changepoint detection. Following segmentation, the defined segments were scored for strand asymmetry, taking into account the sequence composition of the segment. For example in tumours with T→N/A→N informative mutations the number of Ts on the forward strand is the count of Watson sites G_W_ and the number of T→N mutations is *μ_W_* which together give the Watson strand rate *R_W_=μ_W_/G_W_*. The forward strand count of As and mutations from A likewise give the Crick strand rate *R_C_=μ_C_/G_C_*. From these two rates we calculate a relative difference metric, the mutational asymmetry score *S=(R_W_-R_C_)/(R_W_+R_C_)*.

The parameter *S* scales from 1 all Watson (e.g. DEN T→N mutations) through 0 (50:50 T→N:A→N) to −1 for all Crick (e.g. DEN A→N). For the categorical assignment, *S* ≥ 0.3 is Watson strand asymmetric, *S* ≤ −0.3 Crick strand asymmetric and in the range −0.3 < *S* < 0.3 symmetric, though more stringent filtering was applied where noted. Segments containing <20 informative mutations were discarded from subsequent analyses.

To test for oncogenic selection at sites with recurrent mutations, mutational asymmetry segments overlapping the focal mutation were categorised based on their asymmetry score *S*, as above. The test was implemented as a Fisher’s exact test with the 2×2 contingency table comprising the counts chromosomes (two autosomes per cell) stratified by Watson versus Crick asymmetry and the presence of the focal mutation in the tumour. Tumours containing another known driver gene or recurrent mutation within the focal asymmetry segment were discarded from the analysis. We estimated the minimum recurrence of a mutation necessary to reliably detect oncogenic selection through simulation. Biased segregation of chromosomes containing drivers was modelled using the observed median excess of T→N over A→N lesions (23 fold), and random segregation of non-driver containing strands (1:1 ratio). Our model predicted >33 C3H recurrences or >41 CAST recurrences would give 80% power to detect oncogenic selection if present.

### Tumour cellularity estimates

We calculated tumour cellularity as a function of the non-reference read count in autosomal chromosomes *(1-R/d)*2* where *R* is the reference read count at a mutated site and *d* is the total read depth at the site. For each tumour these values were binned in percentiles and the midpoint of the most populated (modal) percentile taken as the estimated cellularity of the tumour. Given the low rate of copy number variation across the DEN induced tumours, no correction was made for copy-number distortion.

### Identifying and filtering reference genome mis-assemblies

Since lesion segregation, mutation asymmetry patterns allow the long-range phasing of chromosome strands, they can detect discrepancies in sequence order and orientation between the sequenced genomes and the reference. We identified autosomal asymmetry segments that immediately transitioned from Watson bias (*S* > 0.3) to Crick (*S* < −0.3) or vice versa without occupying the intermediate unbiased state (−0.3 > *S* < 0.3); such “discordant segments” are unexpected. Allowing for ±100kb uncertainty in the position of each exchange site we produced the discordant segment coverage metric. At sites with discordant segment coverage >1 we calculated percentage consensus for mis-assembly *M=ds/(ds+cs)* where *ds* is the number of discordant segments over the exchange site and *cs* the number of concordant: where either Watson or Crick mutational asymmetry extends at least 1×10^6^ nucleotides on both sides of the exchange site.

### Sister chromatid exchange site analysis

Identified sister chromatid exchange sites were aggregated across tumours from each strain. Exchange sites within 1×10^6^ nt of known and proposed reference genome mis-assembly sites were excluded from analysis. The mid-point between the flanking informative mutations was taken as the reference genome position of the exchange event, and the distance between those flanking mutations as the positional uncertainty of the estimate. To generate null expectations for mutation rate measures, the coordinate of an exchange was projected into the genome of a proxy tumour and the mutation rates and patterns measured from that proxy tumour (repeated 100 times). The permutation of tumour identifiers for the selection of proxy tumours was a shuffle without replacement that preserved the total number of exchange sites measured in each tumour.

The comparison of mutation spectra between windows was calculated as the cosine distance between the 96 category trinucleotide context mutation spectra for the whole genome and that calculated for the aggregated 5kb window. The 96 categories were equally weighted for this comparison.

Exchange site enrichment analysis used Bedtools ^53^ shuffle to permute the genomic positions of exchange sites into the analysable fraction of the genome (defined above). Observed rates of annotation overlap were compared to the distribution of values from 1,000 permuted exchange sites. For genic overlaps we used Ensembl v91 ^47^ coordinates for genic spans; gene expression status was based on the summed expression over all annotated transcripts for the gene from P15 liver from the matched mouse strain. Expression thresholds were defined as >50th centile for active and <50th centile for inactive genes.

### Transcription coupled repair calculations

For each protein coding gene, the maximally expressed transcript isoform was identified from P15 liver in the matched strain (TPM expression), subsequently the primary transcripts. In the case of ties, transcript selection was arbitrary. Genes were partitioned into five categories based on the expression of the primary transcript: expression level 0 (<0.0001 TPM) and four quartiles of detected expression.

Using the segmental asymmetry patterns of each tumour and the annotated coordinates (Ensembl v91) of the selected transcripts, we identified transcripts completely contained in a single Watson or Crick asymmetric segment and located at least 200kb from the segment boundary at both ends. We also applied strict asymmetry criteria of mutational asymmetry scores *S* > 0.8 for Watson and *S* < −0.8 for Crick asymmetry segments, though analysis with the standard asymmetry thresholds and no segment boundary margin give similar results and identical conclusions. For each transcript in each tumour we then used both the transcriptional orientation of the gene and the mutational asymmetry of the segment containing it to resolve the segregated lesions to either the template (anti-sense) or non-template (sense) strand of the gene. Transcripts contained in mutationally symmetric regions or not meeting the strict filtering criteria were excluded from analysis.

We then analysed mutation rates stratifying by gene expression level and the template/non-template strand of the lesions but aggregating between tumours within the same strain. The transcription start site coordinates used correspond to the annotated 5’ end of the primary transcripts.

### Multi-allelic variation

Aligned reads spanning genomic positions of somatic mutations were genotyped using Samtools mpileup (v.1.9) ^56^. Genotypes supported by ≥2 reads with a nucleotide quality score of ≥20 were reported, considering sites with two alleles as biallelic, those with three or four alleles as multi-allelic. The fraction of called mutations exhibiting multi-allelic variation was calculated for the analysable fraction of the genome, across 10Mb consecutive windows and also for each of the mutational asymmetry segments calculated for each tumour.

A null expectation for the multi-allelic rate estimate was generated per C3H tumour; genomic positions identified as mutated across the other 370 tumours were down-sampled to match the mutation count in the focal tumour. Any of these proxy mutation sites with a non-reference genotype supported by ≥2 reads and nucleotide quality score ≥20 at the focal site were referred to as “multi-allelic” for the purposes of defining a background expectation for the calling of multi-allelic variation. For each tumour, this was repeated 100 times and the mean reported.

We used whole exome sequencing (WES) of fifteen C3H tumours from ^10^) that have subsequently been used to generate WGS data in this study as a basis for validating multi-allelic calls. Multi-allelic variant positions derived from WGS were genotyped in WES using Samtools mpileup, as described above. Only sites with ≥30x WES coverage were considered and alleles were found to be concordant if a WGS genotype was supported by ≥1 read in the WES data. To provide a null expectation, the analysis was repeated using WES data from a different tumour and validation rates reported for all versus all combinations of mismatched WGS-WES pairs (15^2^-15=210).

To quantify combinatorial genetic diversity for each tumour, pairs of mutations located between 3-150nt apart were phased using sequencing reads that traversed both mutation sites. Distinct allelic combinations were counted after extraction with Samtools mpileup using only reads with nucleotide quality score ≥20 over both mutation sites.

### Estimating the cell generation of transformation

Knowing the faction of lesion segregation segments that generated multi-allelic variation across a tumour genome allows the inference of the generation time post-mutagenesis of the cell from which the tumour developed, because each successive cell generation is expected to retain only 50% of the lesion containing segments. We estimate this fraction as follows.

Let *p* denote the fraction of multi-allelic segments and let q be its complement, i.e. the fraction of non-multi-allelic segments, for each tumour genome. Segment boundaries being sister chromatid exchange sites or chromosome boundaries. In order to determine *p*, we re-purpose the quadratic Hardy-Weinberg equation: *p+q=p^2^+2pq+q^2^ =1*, which holds since the two possible fractions need to sum to unity. Given an asymmetric segment of interest in the diploid genome, there are 3 distinct scenarios: (i) both chromosomes are multi-allelic (*p^2^*), (ii) One of the chromosomes is multi-allelic and the other is not (*pq+qp*) and (iii) both chromosomes are non-multi-allelic (*q^2^*). The first two scenarios are not distinguishable from the data as both appear multi-allelic (*m*). However, in the third scenario, for a segment to be non-multi-allelic (biallelic, *b*), both chromosomal copies have to be non-multi-allelic. As described below, *q^2^* can be estimated directly from the data and is subsequently used to estimate *p=1-sqrt(q^2^)* and hence the cell generation number of transformation post-mutagenesis.

The estimation of *q^2^* requires computing the ratio *q^2^*=*b/(b+m).* We can directly observe the counts of *b* as non-multi-allelic segments. The number of autosomal chromosome pairs (*n*=19) and count of sister chromatid exchange events (*x*) give the total number of segments in the genome *b+m=n+x*. Exchange events are not expected to align between allelic chromosomes which will result in the partial overlap of segments between allelic copies. Although this increases the number of observed segments (*b* and *m*) relative to actual segments, assuming the independent behaviour of allelic chromosomes and that segment length is independent of multi-allelic state, this partial overlap does not systematically distort the quantification of *b* or the estimation of *q^2^*.

To call a non-multi-allelic segment (*b*) we require less than 4% multi-allelic sites. The threshold based on the tri-modal frequency distribution of multi-allelic rates per-segment, aggregated over all 371 C3H tumours. The 4% threshold separates the lower distribution of multi-allelic rates from the mid and higher distributions.

To test for the enrichment of specific driver gene mutations in early generation versus late generation transformation post-DEN treatment, we applied Fisher’s exact test (fisher.test function in R) to compare the generation 1 ratio of tumours with, versus those without, a focal mutation to the same ratio for tumours inferred to have transformed in a later generation. We report the same odds ratios calculated requiring that the “with focal mutation” tumours had a driver mutation in only one of the driver genes: *Hras, Braf*, or *Egfr*.

### Cell-line and human cancer mutation analysis

Somatic mutation calls were obtained from DNA maintenance and repair pathway perturbed human cells ^30^. Of the 128,054 reported single nucleotide variants, 6,587 unique mutations (genomic site and specific change) were shared between two or more sister clones, so likely represent mutations present but not detected in the parental clone. All occurrences of the shared mutations were filtered out leaving 106,688 mutations for analysis, although the inclusion of these filtered mutations does not alter any conclusions drawn. Somatic mutation calls from mutagen exposed cells ^5^ were obtained, no additional filtering was applied to these sub-clone mutations.

Somatic mutation calls from the International Cancer Genome Consortium (ICGC) ^57^ were obtained as simple_somatic_mutation.open.* files from release 28 of the consortium, one file for each project. These somatic mutations have been called from a mixture of whole genome and whole exome sequencing. Of the 18,965 patients represented (and not embargoed in the release 28 dataset), 116 were excluded from analysis; these represent a distinct whole exome sequenced subset of the LICA-CN project that appear to show a processing artefact in the distribution of specific mutation subsets. ICGC mutations were filtered to remove insertion and deletion mutations and also filtered for redundancy so that each mutation was only reported once for each patient.

### The rl_20_ metric and runs tests

Amongst only the informative mutations (e.g. T→N/A→N in DEN) three consecutive T→N without an intervening A→N is a run of three. The R function rle was used to encode the run-lengths for binary vectors of informative mutations along the genome of a focal tumour. Ranking them from the longest to the shortest run, we find the set of longest runs that encompass 20% of all informative mutations in the tumour. The run-length of the shortest of those is reported as the rl_20_ metric. The threshold percent of mutations was defined as having to be less than 50%, as on average only 50% of the autosomal genomes are expected to show mutational asymmetry patterns. On testing with randomised data, the value of 20% gave a stable null expectation (maximum observed value of a run of five) and still encompassed a large fraction of the informative mutations. All rl_20_ results reported were implemented so that runs were broken when crossing chromosome boundaries.

The Wald-Wolfowitz runs test was performed using the runs.test function of the R randtests (v.1.0) ^58^ library. It was applied to binary vectors of informative changes as described above, with threshold=0.5.

The Wald-Wolfowitz runs test significance is inflated by coordinated dinucleotide changes, such as those produced by UV light exposure and also other local mutational asymmetries such as replication asymmetry ^9^ and kataegis events ^18, 59^. The rl_20_ metric appears robust to most such distortions but we find it efficiently detects kataegis events that are in an otherwise mutationally quiet background, as is often the case for breast cancer. For this reason we also indicate the total genomic span of mutations in the rl_20_ subset of mutation runs: kataegis events typically span a tiny (<5%) fraction of the whole genome.

### Computational analysis environment

Primary data processing was performed in shell-scripted environments calling the software indicated. Except where otherwise noted, analysis processing post-variant calling was performed in a Conda environment and choreographed with Snakemake running in an LSF batch control system (**Supplementary Table 3**).

### Code and data availability

The analysis pipeline including Conda and Snakemake configuration files can be obtained from the repository https://git.ecdf.ed.ac.uk/taylor-lab/lce-ls. The WGS BAM files are available from the European Nucleotide Archive (ENA) under accession: PRJEB15138. RNA-seq files are available from Array Express E-MTAB-8518. Digitised histology images are available from Biostudies under accession S-BSST129.

### Key resources

The key reagents and resources required to replicate our study are listed in **Supplementary Table 3**. For externally sourced data, where applicable, URLs that we used can be found in the Git repository https://git.ecdf.ed.ac.uk/taylor-lab/lce-ls.

## Acknowledgements and funding

We gratefully acknowledge Maša Roller and Florian Markowetz for supervision, and Loris Mularoni and Graham Ritchie for software support. Also, the valuable contributions by the CRUK Cambridge Institute Core facilities: CRUK Biological Resources (Angela Mowbray), Preclinical Genome Editing (Lisa Young, Steven Kupczak, Maureen Cronshaw, Paul Mackin, Yi Cheng, Lena Hughes-Hallett), Genomics (James Hadfield, Fatimah Bowater), Bioinformatics (Gord Brown, Matthew Eldridge, Richard Bowers), Histopathology and ISH, Research Instrumentation, and Biorepository; Edinburgh Genomics (Clinical) Facility; and the EMBL-EBI technical services cluster (Zander Mears, Andrea Cristofori, Tomasz Nowak, Sundeep Nanuwa, Vahit Tabak, Alessio Checcucci).

This work was supported by: Cancer Research UK (20412, 22398), the European Research Council (615584, 682398), the Wellcome Trust (WT108749/Z/15/Z, WT106563/Z/14/A, WT202878/B/16/Z), the European Molecular Biology Laboratory, the MRC Human Genetics Unit core funding programme grants (MC_UU_00007/11, MC_UU_00007/16), the ERDF/Spanish Ministry of Science, Innovation and Universities-Spanish State Research Agency/DamReMap Project (RTI2018-094095-B-I00). S.J.A. received a Wellcome Trust PhD Training Fellowship for Clinicians (WT106563/Z/14/Z) and is now funded by is funded by a National Institute for Health Research (NIHR) Clinical Lectureship. O.P. is funded by a BIST PhD fellowship supported by the Secretariat for Universities and Research of the Ministry of Business and Knowledge of the Government of Catalonia and the Barcelona Institute of Science and Technology. V.S. is supported by an EMBL Interdisciplinary Postdoc (EIPOD) fellowship under Marie Skłodowska-Curie actions COFUND (664726). E.K. is supported by the EMBL International PhD Programme. C.A-P. is supported by La Caixa Foundation fellowship ((ID 100010434; LCF/BQ/ES18/11670011). A.E. is supported by a UKRI Innovation Fellowship (MR/RO26017/1). A.K. is a cross-disciplinary postdoctoral fellow supported by funding from the University of Edinburgh and Medical Research Council (core grant to the MRC Institute of Genetics and Molecular Medicine).

## Author contributions

S.J.A., F.C., C.F., D.T.O. conceived the project and designed the experiments. S.J.A., F.C., C.F., performed the mutagenesis experiments and sequencing experiments. E.L-A, A.M.R. performed supporting experiments. J.S-L provided contract sequencing. S.J.A. performed the histopathological analyses with advice from S.E.D.. C.J.A., M.S.T. performed computational analysis. M.S.T. discovered lesion segregation. O.P., V.S., T.F.R., M.L., S.A., E.K., J.L. performed supporting computational analysis. C.A-P., S.B., R.D., A.E., V.B.K., A.K., I.S., L.T. contributed to the computational analyses. T.F.R., M.L., S.A., A.D.Y. curated data. S.J.A., C.A.S., N.L.B., P.F., D.T.O., M.S.T. supervised the work. S.J.A., C.A.S., N.L.B., P.F., D.T.O., M.S.T. lead the Liver Cancer Evolution Consortium. S.J.A. and P.F. provided scientific and administrative organisation. S.J.A., C.A.S., N.L.B., P.F., D.T.O., M.S.T. funded the work. S.J.A., D.T.O., M.S.T. wrote the manuscript. All authors had the opportunity to edit the manuscript. All authors approved the final manuscript.

## Extended data

**Extended Data Fig. 1.**
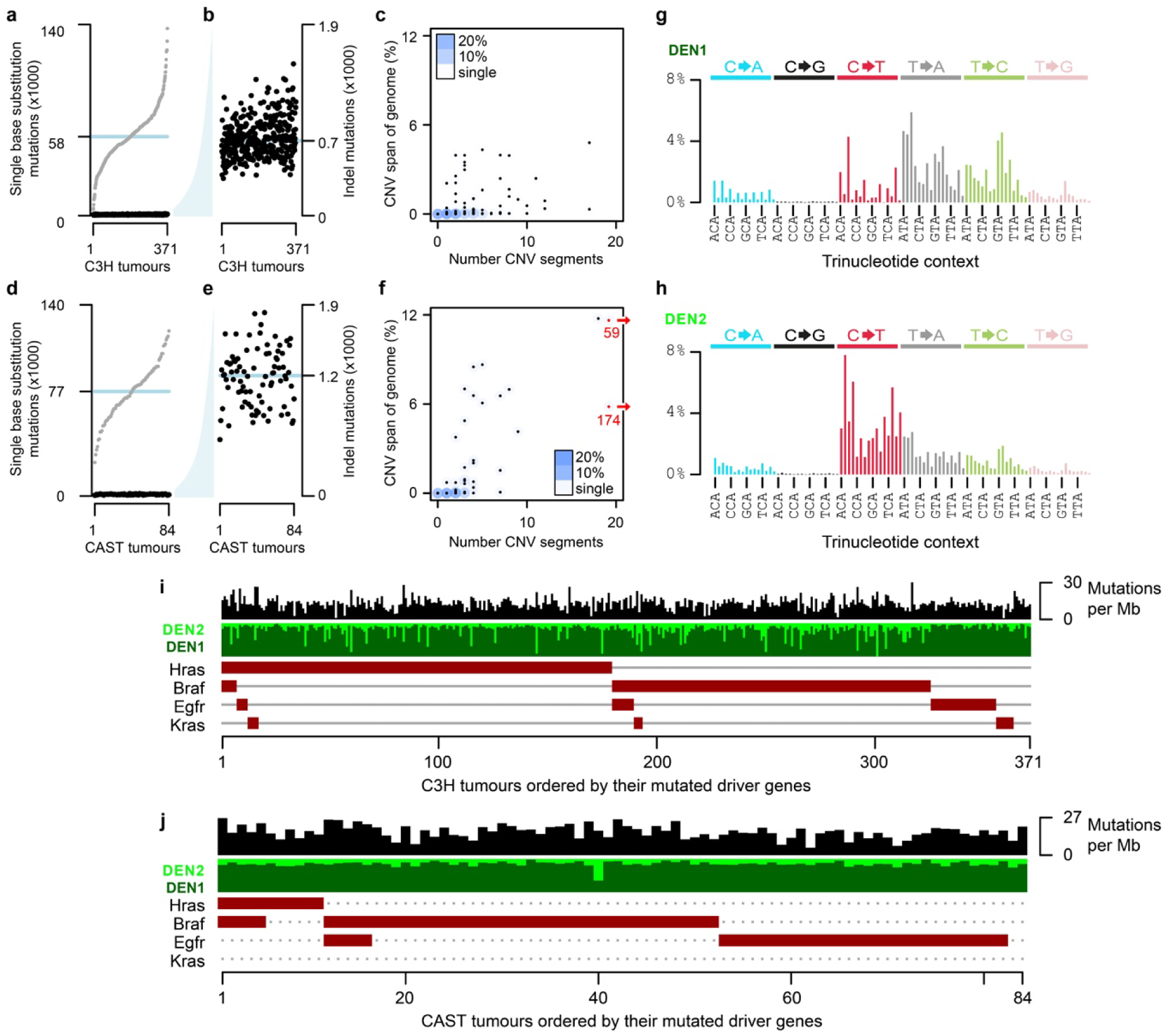
Summary mutation metrics for both C3H and CAST tumours. **a**, Single nucleotide substitution rates per C3H tumour, rank ordered over x-axis (grey points, median blue line). Insertion/deletion (indel, <11 nt) rates show as black. **b**, Y-axis from **a**, expanded to show distribution of indel rates with preserved tumour order. **c**, Number of C3H copy number variant (CNV) segments and their total span as a percent of the haploid genome. Blue shading shows intensity of overlapping points as a percent of all tumours in the plot. **d-f**, Corresponding plots for CAST derived tumours, **f**, two extreme x-axis outliers relocated (red) and x-axis value shown. **g-h**, Mutation spectra deconvolved from the aggregate spectra of 371 C3H tumours, subsequently referred to as the DEN1 and DEN2 signatures. **i**, Oncoplot summarising mutation load, mutation spectra, and driver gene mutation complement of C3H tumours. **j**, Oncoplot of CAST derived tumours as **i**.

**Extended Data Fig. 2.**
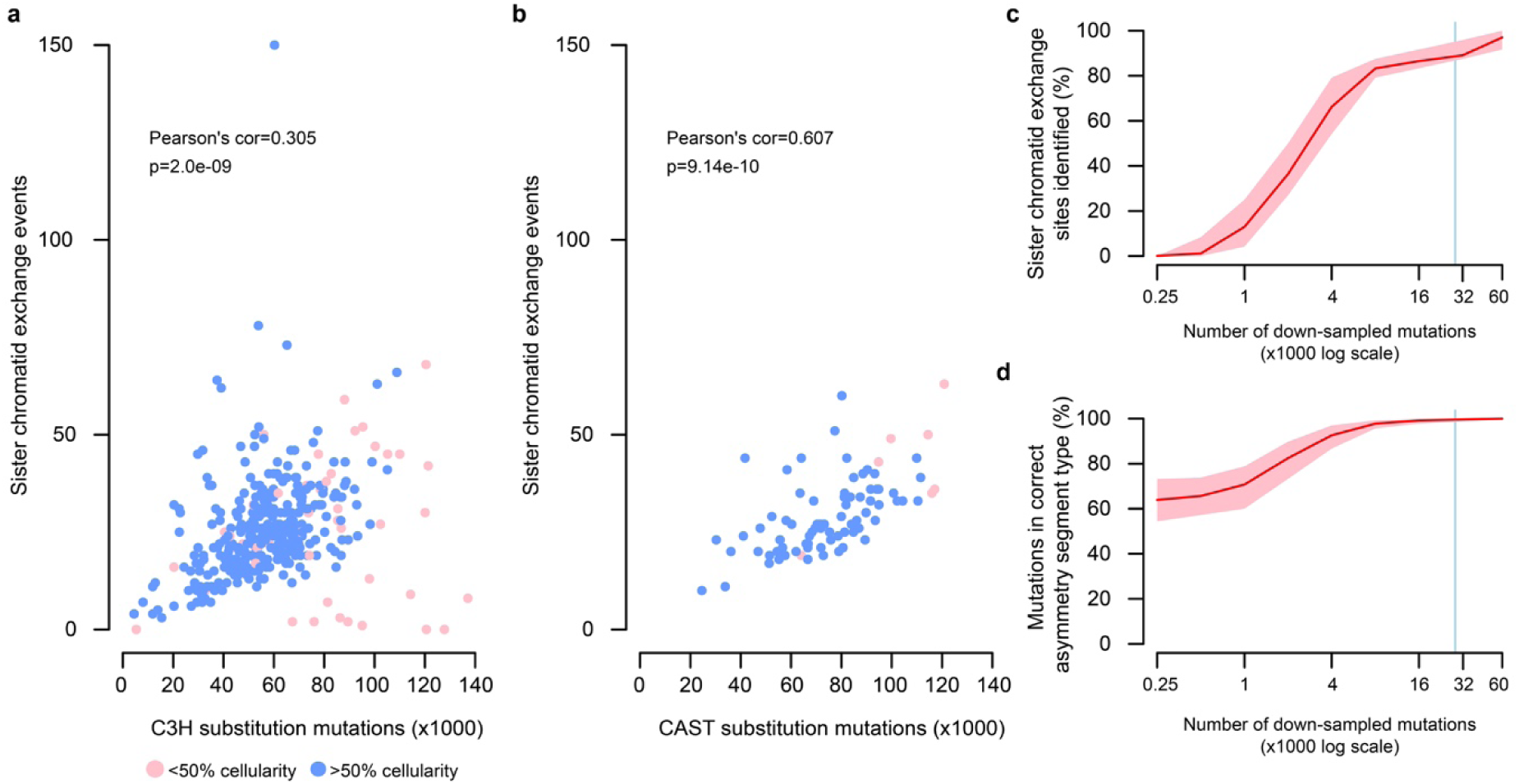
The frequency of sister chromatid exchanges is correlated with the point mutation burden. **a**, The relationship between single nucleotide substitution mutation load and detected sister chromatid exchange events in C3H tumours. Tumours with low cellularity (pink) have high mutation load and form a sub-group with few detected sister chromatid exchange events; these are suspected to be polyclonal tumours. **b**, As for **a** but showing CAST derived tumours. **c**, Evaluation of the relationship between mutation load and ability to detect sister chromatid exchange events. Mutations from C3H tumour 94315_N8 (shown in Fig. 2) randomly down-sampled and segmentation analysis applied. Y-axis shows the percentage of sister chromatid exchange events detected (100 replicates, 95% C.I. pink). X-axis is on a log-scale: 95% of C3H and >95% of CAST tumours have mutation counts to the right of the blue vertical line. Down-sampling other tumours gave comparable results. **d**, The same down-sampling data as shown in panel **c** but the y-axis shows the percent of mutations with the correct (same as full data) mutational asymmetry assignment.

**Extended Data Fig. 3.**
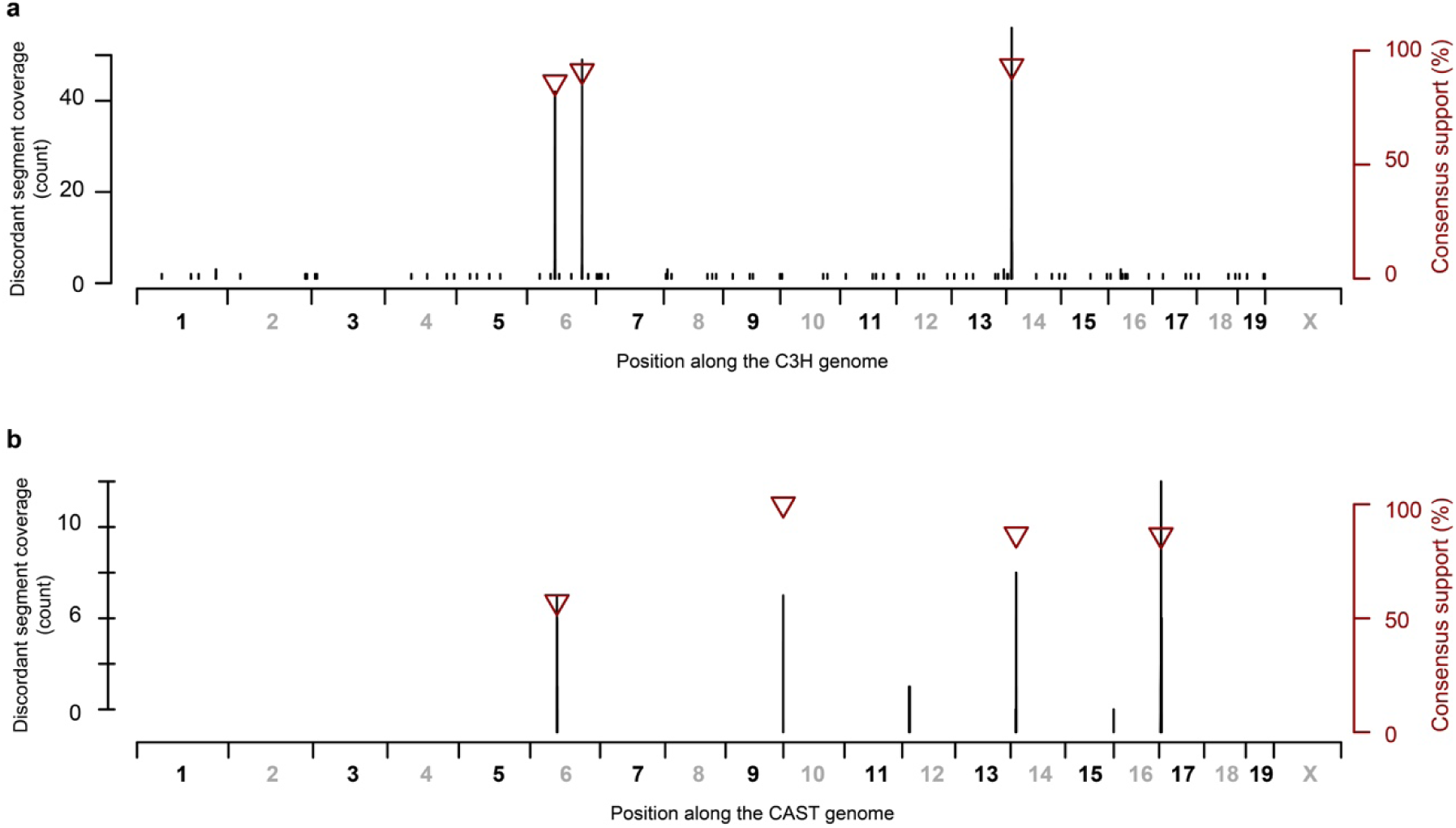
Localising candidate reference genome assembly errors. Genome coordinates shown on the x-axis. Separate plots are shown for the C3H (**a**) and CAST (**b**) strains. Immediate switches between Watson and Crick mutational strand asymmetry are not expected on autosomes unless both copies of the chromosome have a sister chromatid exchange event at equivalent sites. However, inversions and inter-chromosomal translocations between the sequenced genomes and the reference assembly are expected to produce immediate asymmetry switches. Recurrent evidence of such a site indicates a systematic discrepancy between the sequence of the reference genome and the actual genomic DNA sequence. The discordant segment coverage (DSC) count (black y-axis) shows the number of informative tumours (those with either Watson or Crick strand asymmetry at the corresponding genome position) that suggest a tumour genome to reference genome discrepancy at the indicated genomic position. The consensus support (brown y-axis) plotted as triangles shows the percentage of informative tumours that support a genomic discrepancy at the indicated position (only shown for values >50% support). The two sites on chromosome 6 in C3H correspond to a previously identified C3H strain specific inversion that is known to be incorrectly oriented in the C3H reference assembly^60^. Similarly we detect four candidate mis-assemblies in the CAST reference genome, though these could represent mouse colony specific chromosomal rearrangements. The candidate mis-assembly on C3H chromosome 14 is at an approximately orthologous position to the chromosome 14 site in CAST suggesting that this may represent a rearrangement shared between strains or a missassembly in the BL6 GRCm38 reference assembly against which other mouse reference genome assemblies have been scaffolded.

**Extended Data Fig. 4.**
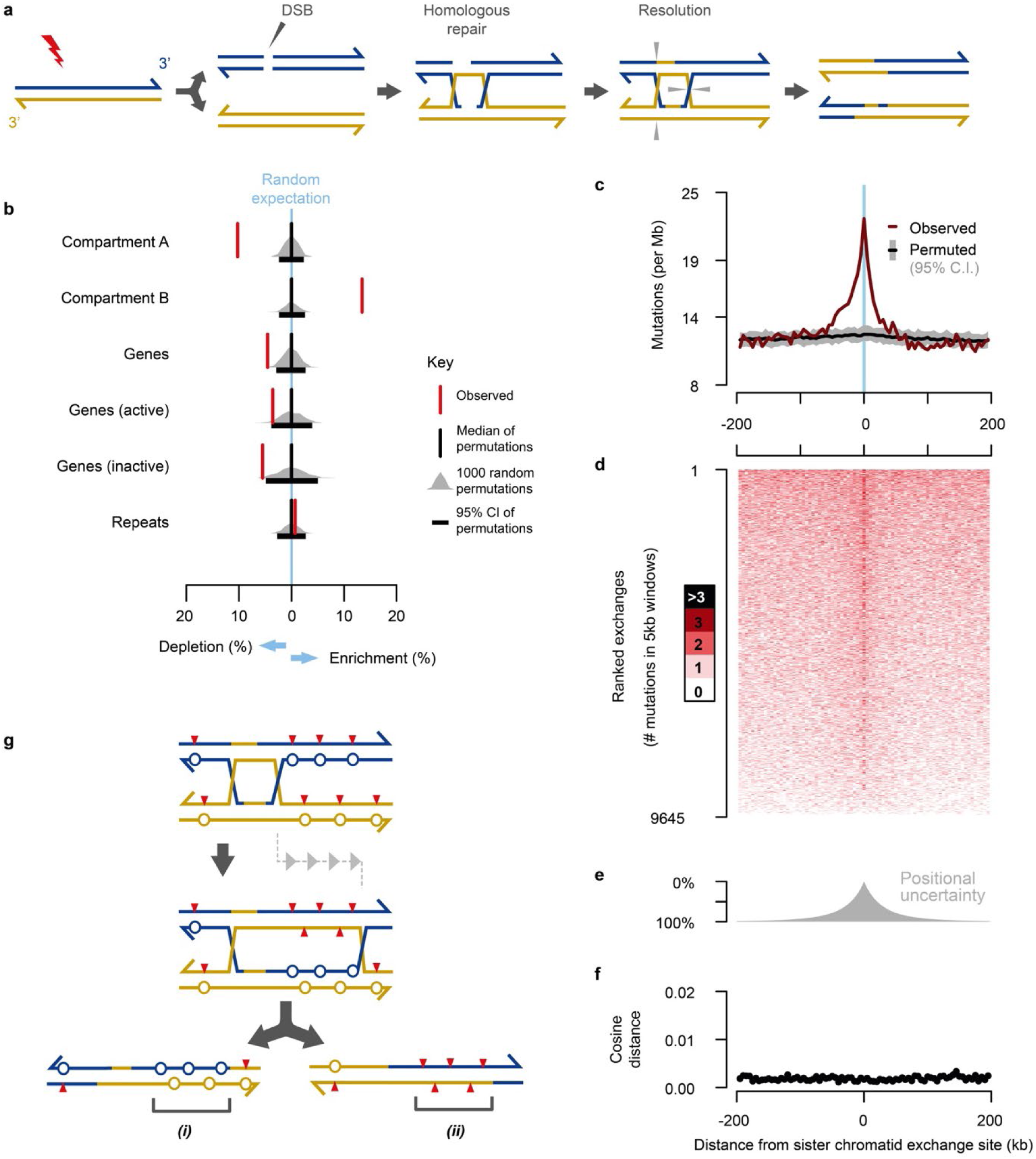
Locally elevated mutational diversity is driven by sister-chromatid exchange. **a**, Double strand breaks (DSBs) and other DNA damage can trigger homologous recombination (HR) mediated DNA repair between sister chromatids. The repair intermediate resolves into separate chromatids through cleavage and ligation; grey triangles denote cleavage sites for one of the possible resolutions that would result in a large-scale sister-chromatid exchange event. **b**, Enrichment analysis of sister chromatid exchanges sites (red) compared with null expectations from randomly permuting locations into the analysable fraction of the genome (grey distributions), the black boxes denote 95% of 1,000 permutations. Sister chromatid exchange events are enriched in later replicating and transcriptionally less active genomic regions (Hi-C defined compartment B), and correspondingly depleted from early replicating active regions. **c**, Aggregating across n=9,645 sister chromatid exchange sites, the observed mutation rate approximately doubles at the inferred site of exchange (x=0). Aggregate mutation rates (brown) were calculated in consecutive 5kb windows. Compositionally matched null expectation was generated by permuting each exchange site into 100 proxy tumours and calculating median (black) and 95% confidence intervals (grey) while preserving the total number of projected sites per proxy tumour. **d**, The elevated mutation count is not the result of a high mutation density in a subset of exchange sites, rather it is a subtle increase in mutations across most exchange sites. Heatmap showing mutation counts calculated in consecutive 5kb windows across each exchange site. Rows represent each exchange site, rank-ordered by total mutation count across each 400kb interval. **e**, The distribution of positional uncertainty in exchange site location approximately mirrors the decay profile of elevated mutation frequency. **f**, Divergence of mutation rate spectra is shown as cosine distance between the analysed window and the genome wide mutation rate spectrum aggregated over all C3H tumours. Despite the elevated mutation frequency, there is no detected distortion of the mutation spectrum. **g**, A model based on HR repair intermediate, branch migration that produces heteroduplex segments of *(i)* mismatch:mismatch (circles) and *(ii)* lesion:lesion (red triangles) strands. Subsequent strand segregation would increase the mutational diversity of a descendant cell population but not the mutation count per cell (key as per Fig. 2).

**Extended Data Table 1.** A lesion segregation based test for oncogenic selection.

| Strain | Gene | Mutation | Mutation count | Odds ratio | P-value | Known driver |
| --- | --- | --- | --- | --- | --- | --- |
| C3H | Braf | 6:37548568_A/T | 151 | 2.13 | 5.77x10<sup>-6</sup> | Yes |
| C3H | Hras | 7:145859242_T/C | 81 | 2.67 | 6.88x10<sup>-6</sup> | Yes |
| C3H | Hras | 7:145859242_T/A | 65 | 1.02 | 1 | Yes |
| C3H | Intronic Fmnl1 | 11:105081902_A/C | 44 | 1.03 | 1 | No |
| C3H | Intergenic | 9:73125689_G/C | 42 | 1.13 | 1 | No |
| C3H | Egfr | 11:14185624_T/A | 34 | 3.87 | 1.23x10<sup>-4</sup> | Yes |
| CAST | Braf | 6:37451282_A/T | 42 | 1.41 | 0.338 | Yes |

Recurrently mutated sites in both C3H and CAST with sufficient estimated power to detect oncogenic selection through biased strand retention analysis (required >33 C3H recurrences or >41 CAST recurrences). Odds ratio values >1 indicate the predicted correlation of driver mutation and Watson/Crick strand retention in tumours with the candidate driver mutation, but not for those without the mutation. The Fisher's exact test P-value is shown after Bonferroni correction. Known driver indicates the mutation or its orthologous change has previously been implicated as a driver of hepatocellular carcinoma<sup>10</sup>. The CAST 6:37451282\_A/T mutation is orthologous to the C3H 6:37548568\_A/T mutation.

**Extended Data Fig. 5.**
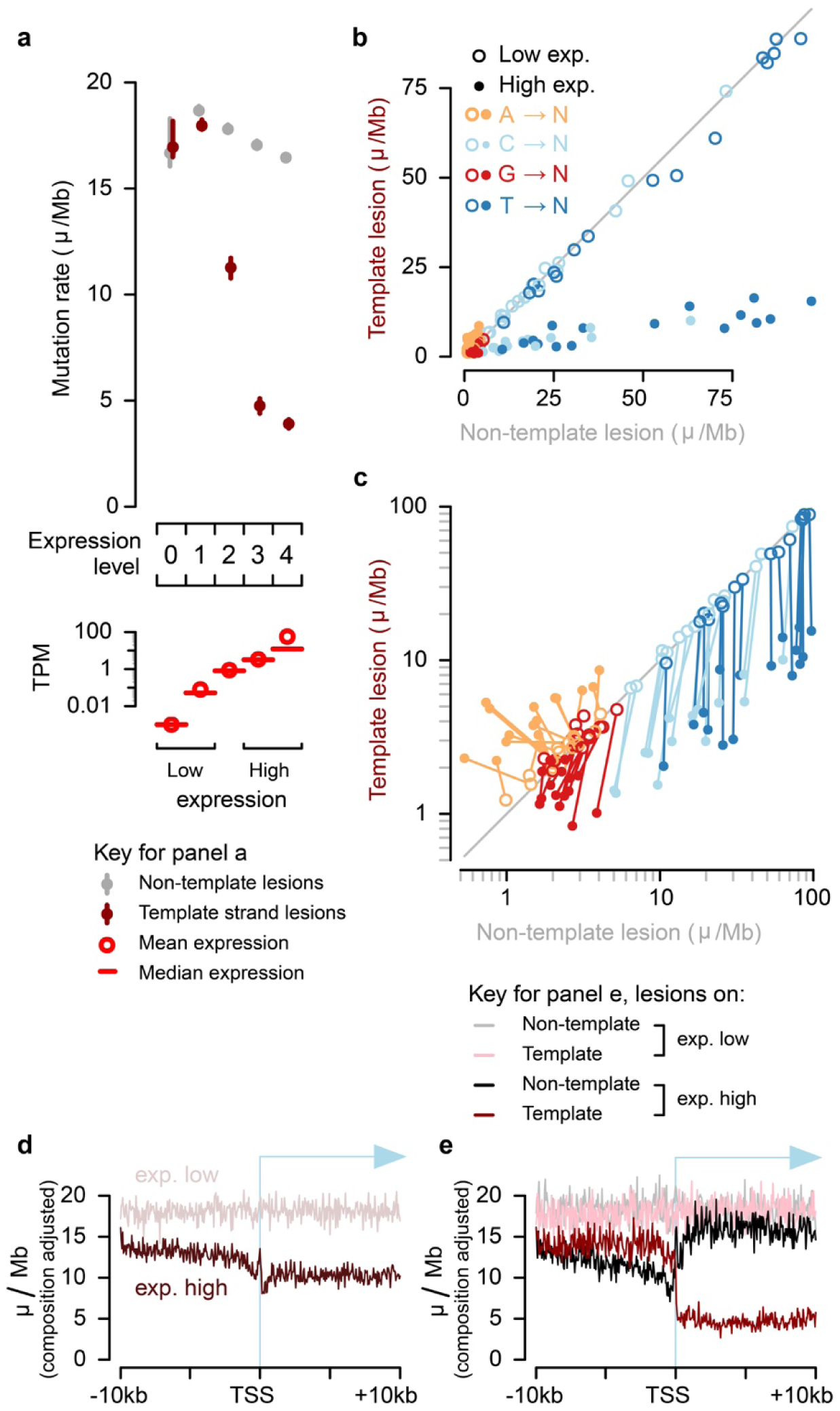
Replication of transcription coupled repair with lesion strand resolution in *Mus castaneus*. **a**, Transcription coupled repair of template strand lesions is dependent on transcription level (P15 liver, transcripts per million (TPM)). Confidence intervals (99%) are shown as whiskers, where broad enough to be visible. **b**, Comparison of mutation rates for the 64 trinucleotide contexts: each context has one point for low and one point for high expression. **c**, Data as in panel **b** plotted on log scale; there is a line linking low and high expression for the same trinucleotide context. **d**, Sequence composition normalised profiles of mutation rate around transcription start sites (TSS). **e**, Stratifying the data plotted in **d** by lesion strand reveals much greater detail on the observed mutation patterns, including the pronounced influence of bidirectional transcription initiation.

**Extended Data Fig. 6.**
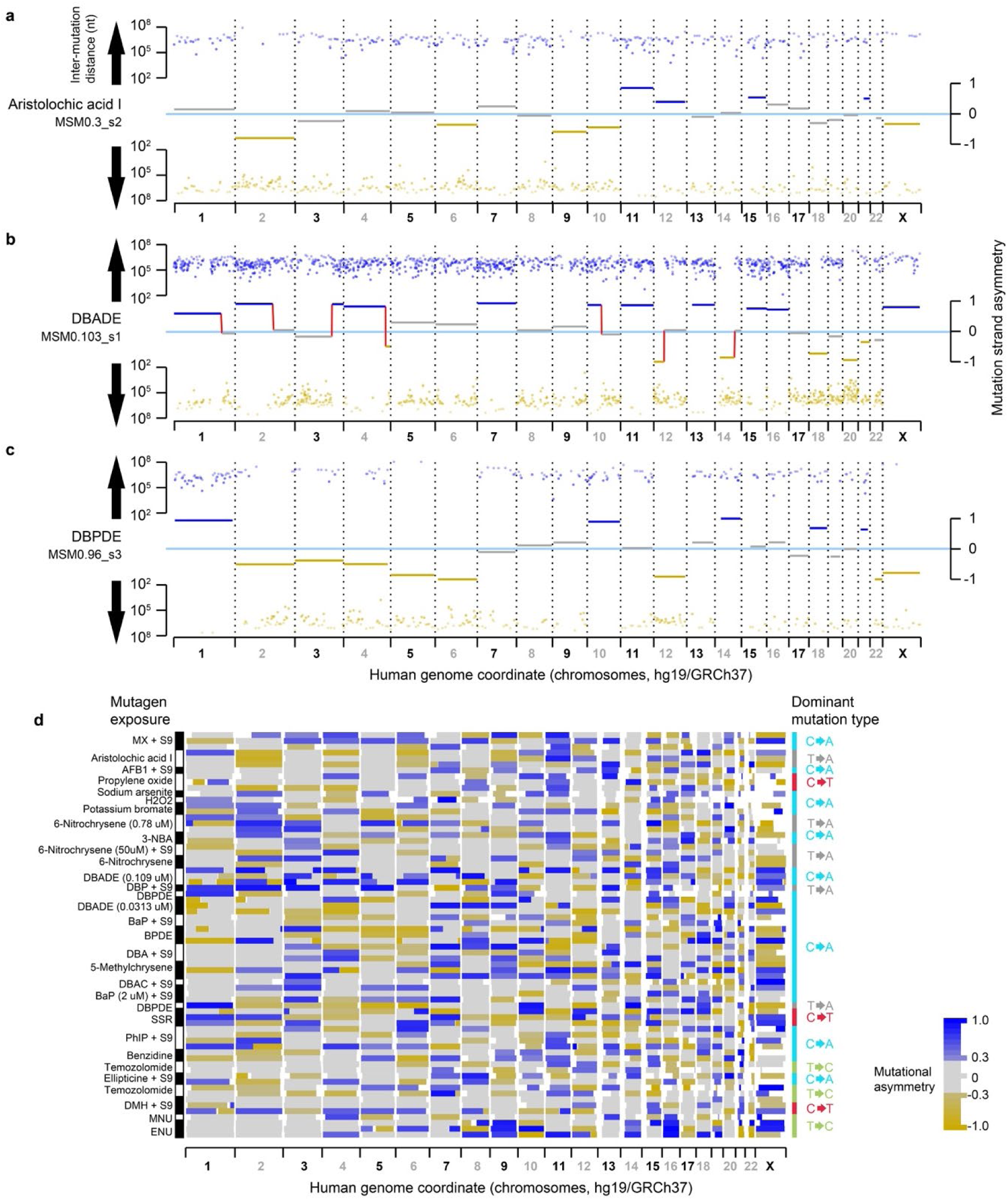
Additional examples of mutation patterns generated by lesion segregation from a diverse range of clinically relevant mutagens. **a**, Despite the low total mutation load (1,308 nucleotide substitutions, 842 informative T→A changes), the mutational asymmetry of lesion segregation (plotted as per Fig. 2a-c) is evident for aristolochic acid exposed clone MSM0.3_s2^5^. **b,c**, Equivalent asymmetry plots for other mutagenised, human induced pluripotent stem cells. **d**, Summary mutation asymmetry ribbons (as per Fig. 2d) for all mutagen exposed clones with rl_20_ >5, which illustrates the independence of asymmetry pattern between replicate clones, almost universal asymmetry on chromosome X, and approximately 50% of the autosomal genome with asymmetry over autosomal chromosomes. The dominant mutation type is indicated for each mutagen. In those clones with low mutation rates, some sister exchange sites are likely to have been missed leading to reduced asymmetry signal (e.g. on the X chromosome, **Extended Data** Fig. 2c).

**Extended Data Fig. 7.**
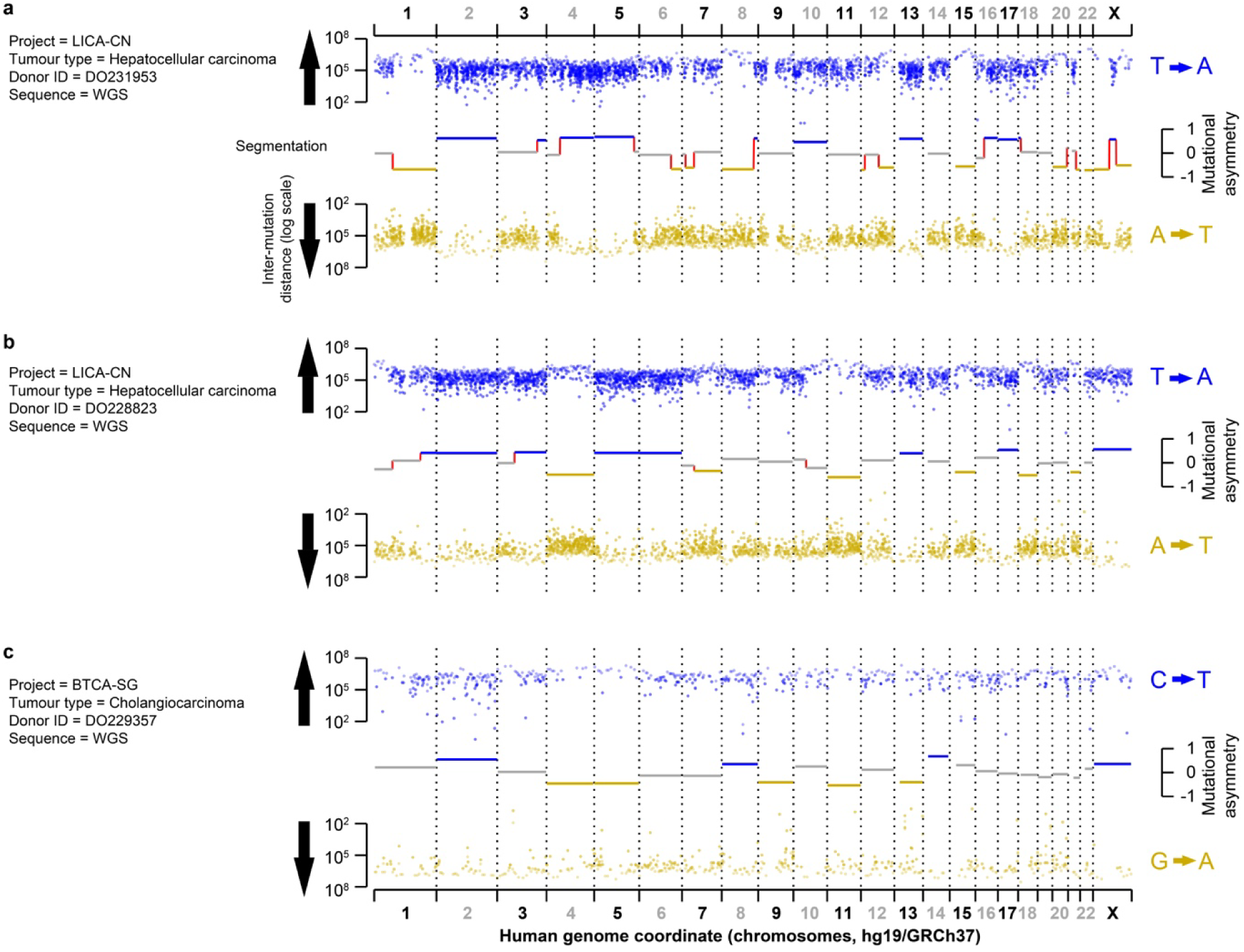
Example mutation asymmetry plots for human cancers with evidence of lesion segregation patterns. **a,b** Hepatocellular carcinomas arising in male patients (project LICA-CN) showing prevalent T→A/A→T substitutions, consistent with aristolochic acid exposure, and mutation strand asymmetry (plotted as per Fig. 2a**-c**). **c**, Cholangiocarcinoma with a lower total mutation load than a,b which shows chromosome scale mutational asymmetry and a distinct mutation signature, dominated by C→T/G→A substitutions of unknown aetiology (plotted as per Fig. 2a**-c**).

**Supplementary Table 1 | Table of tumours sequenced containing key parameters & mutation spectra signature matrices** (Excel file).

**Supplementary Table 2 | Table of Exogenous mutagen and ICGC scan results** (Excel file).

**Supplementary Table 3 | Table of key resources and software** (Excel file).

